# Transcriptome-wide interrogation of the functional intronome by spliceosome profiling

**DOI:** 10.1101/226894

**Authors:** Weijun Chen, Jill Moore, Hakan Ozadam, Hennady P. Shulha, Nicholas Rhind, Zhiping Weng, Melissa J. Moore

## Abstract

Full understanding of eukaryotic transcriptomes and how they respond to different conditions requires deep knowledge of all sites of intron excision. Although RNA-Seq provides much of this information, the low abundance of many spliced transcripts (often due to their rapid cytoplasmic decay) limits the ability of RNA-Seq alone to reveal the full repertoire of spliced species. Here we present “spliceosome profiling”, a strategy based on deep sequencing of RNAs co-purifying with late stage spliceosomes. Spliceosome profiling allows for unambiguous mapping of intron ends to single nucleotide resolution and branchpoint identification at unprecedented depths. Our data reveal hundreds of new introns in *S. pombe* and numerous others that were previously misannotated. By providing a means to directly interrogate sites of spliceosome assembly and catalysis genome-wide, spliceosome profiling promises to transform our understanding of RNA processing in the nucleus much like ribosome profiling has transformed our understanding mRNA translation in the cytoplasm.

## Introduction

Current methods for eukaryotic transcriptome annotation rely heavily on RNA-Seq data from polyA+ or rRNA-depleted cellular RNA wherein intron positions are inferred from reads spanning exon junctions. Exon junction reads are also used to quantify alternative splicing patterns (Trapnell et al., 2012; Wang et al., 2008; 2009). However, due to biases inherent in all RNA-Seq preparation methods (Hu et al., 2014; Li and Jiang, 2012) and any assumptions made during exon junction mapping (e.g., maximum intron length, number of allowed mismatches), not all intron positions can be accurately identified from exon junction reads alone (Tuerk et al., 2017). Further, because individual mRNA and lncRNA isoforms can have widely different decay rates (Clark et al., 2012; Geisberg et al., 2014; Wan et al., 2012), a steady-state snapshot of exon junction reads is unlikely to accurately reflect the flux through different alternative splicing pathways. Finally, if sequencing depth is insufficient and a spliced mRNA is highly unstable (e.g., AS-NMD and snoRNA host gene mRNAs)(McGlincy and Smith, 2008), traditional RNA-Seq may miss the exon-containing spliced product all together.

A more direct approach for mapping intron locations and measuring splicing flux would be to interrogate the introns themselves. One method is to analyze branched reads in RNA-Seq data (Awan et al., 2013; Bitton et al., 2014; Mercer et al., 2015; Taggart et al., 2012). Branches are generated during the first chemical step of splicing wherein the 2’-OH of an adenosine near the 3’ end of the intron (the branchpoint; BP) attacks the 3’-5’ phosphodiester at the 5’ end of the intron (the 5’ splice site; 5’SS) to liberate the 5’ exon terminating with 3’-OH and form a 2’-5’ bond between the BP and the 5’SS nucleotide. Subsequent attack of the 5’ exon on the 3’ end of the intron (the 3’ splice site; 3’SS) during the second step of splicing frees the branched (aka lariat) intron and ligates the two exons. Subsequent debranching of the excised intron by debranching enzyme results in a linear intron beginning with a 5’-phosphate (5’-P) and ending with a 3’-OH.

Branches are incidentally captured during RNA-Seq library construction when reverse transcriptase initiates downstream of the 5’SS and traverses the 2’-5’ bond to continue cDNA synthesis upstream of the BP. However, the low abundance of such inverted reads in traditional RNA-Seq datasets (ca. 1:10^6^; Taggart et al., 2012) limits the utility of this approach. Although circular RNA species can be enriched by 2D electrophoresis (Awan et al., 2013) or exonucleolytic degradation of linear species (Mercer et al., 2015) these approaches work well only for relatively short introns whose lariats are less susceptible to breakage during sample workup. Thus, despite intensive bioinformatics analyses interrogating billions of RNA-Seq reads, <25% of human introns currently have mapped BPs (Mercer et al., 2015; Taggart et al., 2017). In *Schizosaccharomyces pombe*, where the longest known nuclear intron is just 1126 nts, BPs have been validated in only 31% of all introns (Awan et al., 2013; Bitton et al., 2014).

An alternate means to enrich for introns is to purify spliceosomes, the macromolecular machines responsible for identifying splice sites and catalyzing intron excision. Employing a split TAP-tag approach, we previously reported purification, complete proteomics, and preliminary RNA-Seq characterization of an endogenous *S. pombe* spliceosomal complex containing U2, U5 and U6 snRNAs and the Prp19 complex (NTC) (Chen et al., 2014). This endogenous U2·U5·U6·NTC species contains predominantly lariat intron product, so is often referred to as intron lariat spliceosome (ILS) complex (Chen et al., 2014). It was the first complete spliceosome for which a high-resolution structure was determined by cryo-EM (Hang et al., 2015; Yan et al., 2015).

Here we present multiple deep sequencing analyses of the transcripts contained within the *S. pombe* U2·U5·U6·NTC complex. Together, this analysis suite constitutes “spliceosome profiling” (**Figure 1**). Spliceosome profiling includes spliceosome RNA-Seq libraries to investigate overall coverage of introns and exons associated with late stage spliceosomes, 5’-P and 3’-OH libraries of debranched RNA to unambiguously identify sites of spliceosome catalysis at nucleotide resolution, and spliceosome footprint libraries to precisely map the intron regions forming stable interactions with the splicing machinery. These datasets enabled us to annotate hundreds of new introns and correct the annotations of many others, revealing numerous novel mRNA isoforms. We also found evidence that a few *S. pombe* introns are removed by recursive splicing (zero-length exons) and that, unlike group II self-splicing introns, intron excision by the spliceosome requires branch formation. Finally, by developing a new random forest approach to detect BPs in spliceosome RNA-Seq and 3’-OH libraries not subjected to debranching, we were able to identify BPs in 86% of all *S. pombe* introns. Spliceosome profiling thus allows for functional “intronome” (Qin et al., 2016) identification and analysis at unprecedented depths. By combining genome-wide intronome and “exonome” (i.e., “transcriptome”) data, a substantially more comprehensive understanding of post-transcriptional gene regulation can be achieved than is currently obtainable with exonome data alone.

**Figure 1.**
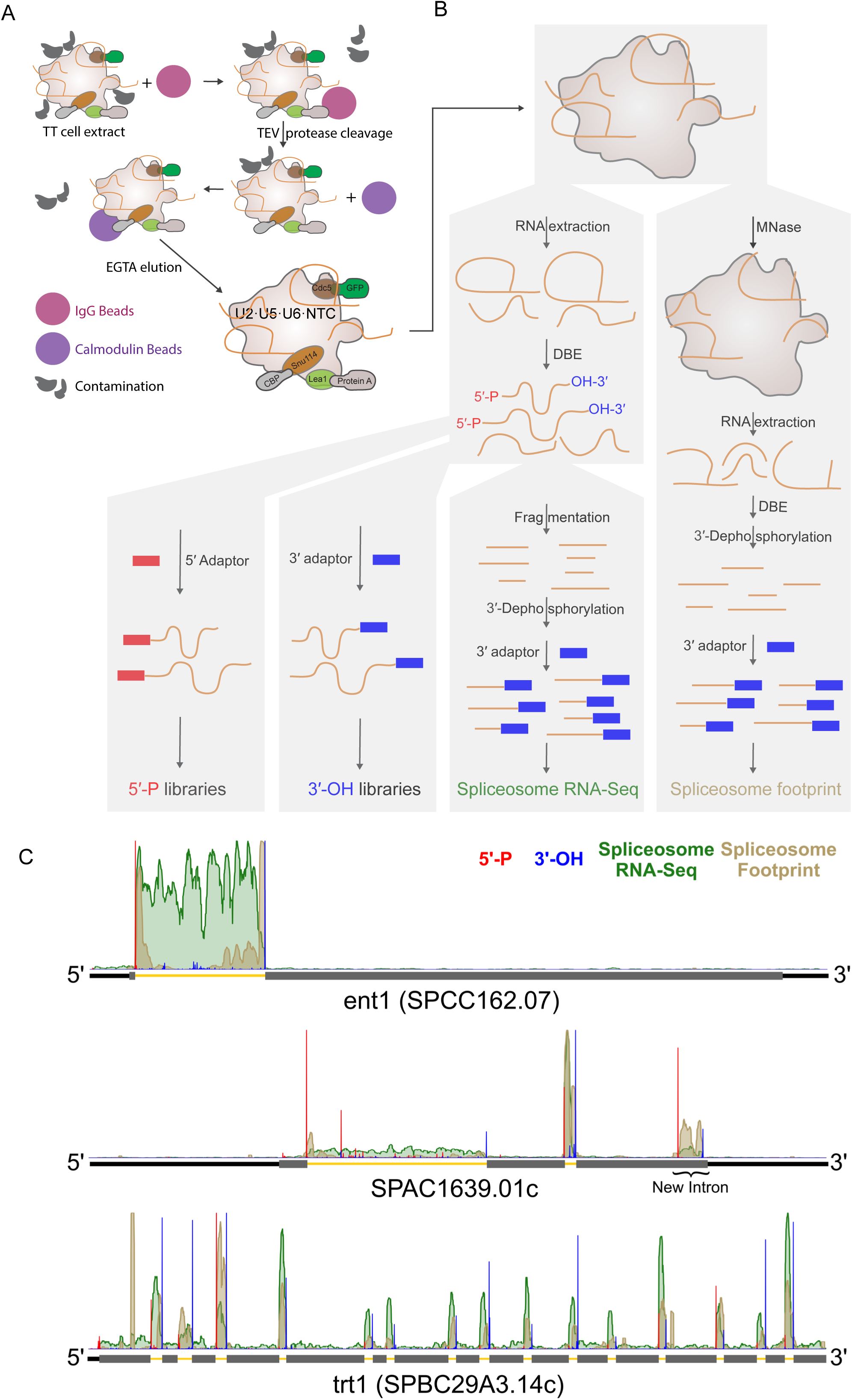
U2·U5·U6·NTC complex purification and library preparation strategies. (A) Scheme of U2·U5·U6·NTC complex purification from triply-tagged (TT) *S. pombe* strain. (B) Scheme of 5’-P, 3’-OH, spliceosome RNA-Seq, and spliceosome footprint library preparation. For 5’-P and 3’-OH libraries, samples were divided following RNA extraction and DBE treatment and used directly for either 5’ or 3’ adaptor ligation. For spliceosome RNA-Seq libraries, extracted RNA was fragmented and 3’ dephosphorylated prior to 3’ adaptor ligation. For spliceosome footprint libraries, unprotected RNA in U2·U5·U6·NTC complexes was digested with micrococcal nuclease (MNase) prior to RNA extraction, 3’ dephosphorylation and 3’ adaptor ligation. (C) Genome browser screenshot of individual genes containing one, two, or fifteen previously-annotated introns and one new intron. Normalized 5’-P (red), 3’-OH (blue), spliceosome RNA-Seq (green) and spliceosome footprint (brown) read coverage was summed across biological replicates. Thick gray rectangles: coding regions; medium black lines: UTRs; thin yellow lines: introns. See also Figure S1 and Table S1

## Results

### U2·U5·U6·NTC complex purification and library characterization

ILS complex should contain a complete record of all sites of spliceosome catalysis. To access this record, we purified U2·U5·U6·NTC complex from two independent batches of haploid *S. pombe* cells grown to late log in rich media. Our tandem affinity purification strategy (Chen et al., 2014) employed a Protein-A tag on the U2 snRNP protein Lea1 (aka, U2 A’) and a calmodulin binding peptide (CPB) on the U5 snRNP protein Snu114 (**Figure 1A**). A GFP tag on the NTC protein Cdc5 allowed for ready monitoring of the purification progress. All three C-terminal tags were inserted in frame with sole chromosomal copy of each protein. Purifications from this triply-tagged (TT) strain were performed under high salt conditions (HS, 700 mM KCl/NaCl) to remove all loosely bound species. As previously described, this approach results in a highly pure preparation of endogenous spliceosomes containing stoichiometric quantities of U2, U5 and U6 snRNAs, core spliceosomal proteins and associated introns (Chen et al., 2014).

Because of the heterogeneous nature of RNAs associated with U2·U5·U6·NTC complex (i.e., snRNAs plus splicing intermediates and products), multiple library types are required to fully profile the contents (**Figure 1B** and **Table S1**). Treatment of purified RNA with recombinant debranching enzyme (DBE) results in linear species amenable to construction of spliceosome RNA-Seq libraries (to yield a complete record of all associated RNAs), 5’-P libraries (to map intron 5’ ends) and 3’-OH libraries (to map intron 3’ ends). Incubation of spliceosomes with micrococcal nuclease (MNase) prior to RNA purification produces spliceosome footprints.

Sequencing on the Illumina HiSeq 4000 and NextSeq platforms yielded 25 to 97 million reads per library, with biological replicates exhibiting correlation coefficients (r) ranging from 0.95 to 0.99 for reads uniquely mapping to annotated genes (*S. pombe* genome version ASM294v2.26; **Table S1A**). Consistent with our previous report that endogenous U2·U5·U6·NTC complex is predominantly ILS complex (Chen et al., 2014), average intron read density was 15-fold higher than average exon read density for both spliceosome RNA-Seq and footprint libraries. This strong enrichment over introns was readily apparent from read coverage on individual genes (**Figure 1C**). In all, 4,499 introns (85% of 5,303 annotated spliceosomal introns) were represented by >10 spliceosome RNA-Seq reads in both biological replicates, with undetected introns generally residing in poorly expressed genes (**Figure S1B**).

The 5’-P and 3’-OH libraries allowed us to precisely map intron ends. As expected, a large fraction of these reads mapped to annotated introns (**Table S1D, E)**, with an aggregation plot revealing striking peaks at intron ends (**Figure 2A and S1C**). In all, 4,691 introns had statistically significant (binomial test, FDR < 0.01) 5’SS 5’-P and 3’SS 3’-OH peaks in both biological replicates. Inclusion of introns with a statistically significant peak at just one end increased this number to 5,105, or 96% of previously annotated *S. pombe* introns. As with the spliceosome RNA-Seq, introns not observed in the 5’-P and 3’-OH libraries corresponded to low abundance mRNAs (**Figure S1B**). Thus our datasets cover the vast majority of intron-containing transcripts expressed during late log growth.

**Figure 2.**
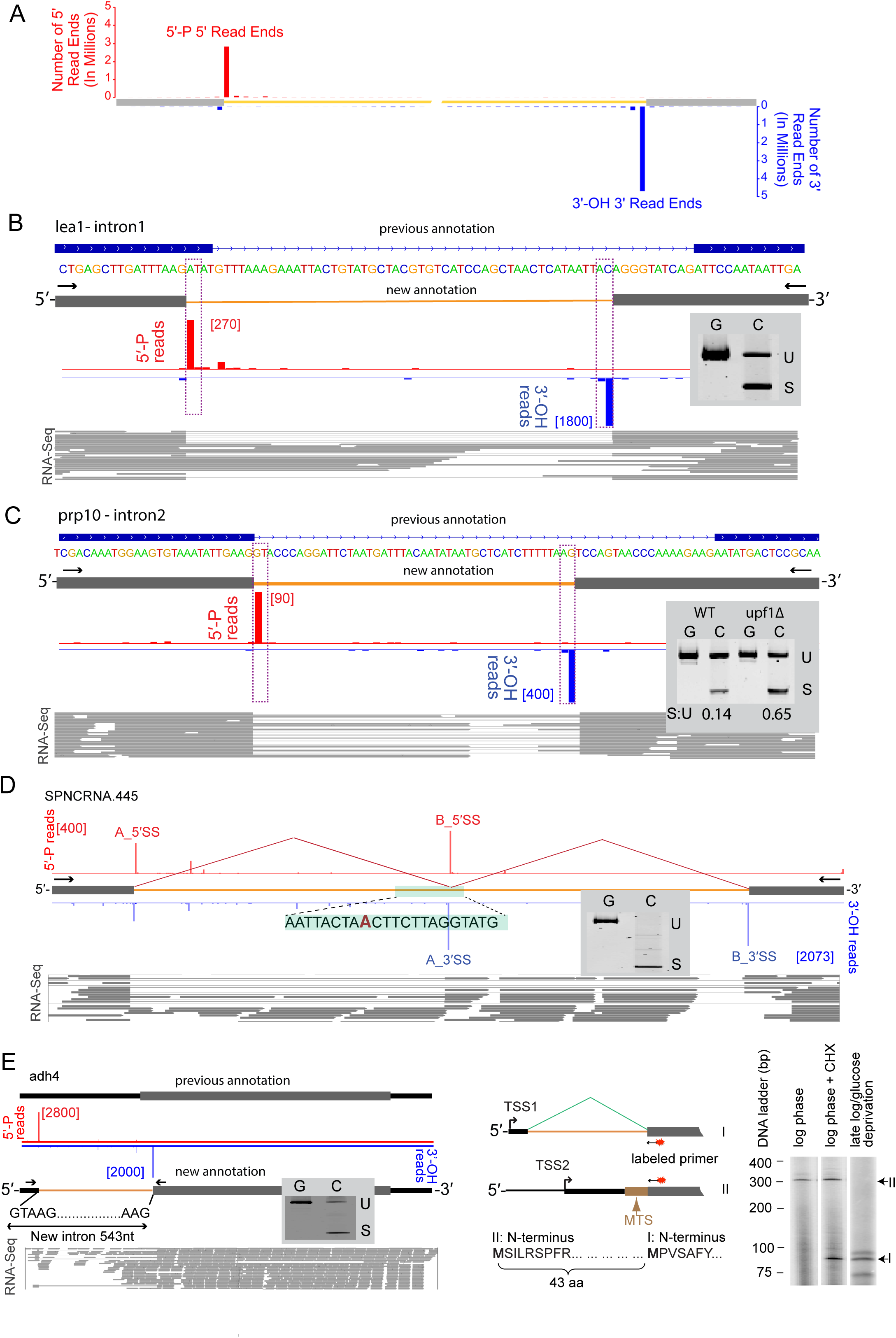
Mapping sites of spliceosome catalysis with 5’-P and 3’ - OH libraries. (A) Aggregation plot of all TT DBE+ 5’-P 5’ read ends (red) and 3’-OH 3’ read ends (blue) across all annotated 5’ and 3’ splice sites in CDS genes. (B) An ATAC intron in lea1 (SPBC1861.08c). Top: previous intron annotation; middle: 5’-P (red) and 3’-OH (blue) read ends from a single biological replicate (read numbers in brackets); bottom: polyA+ WT RNA-Seq reads supporting new annotation. Inset: SYBR gold-stained polyacryamide gel of PCR products (U: unspliced; S: spliced) from genomic DNA (G) or cDNA (C) from WT cells. (C) Corrected intron in prp10 (SPAC27F1.09c). Tracks as in B. Inset: SYBR gold-stained polyacryamide gel of PCR products (U: unspliced; S: spliced) from genomic DNA (G) and cDNA (C) from WT and upf1Δ cells. S : U = ratio of S to U products. (D) Zero-length exon and recursive splicing in SPNCRNA.445 intron. Tracks and inset as in B. (E) A new transcription start site (TSS1) and intron in adh4 gene. Left: Tracks and inset as in B Center: adh4 mRNA isoforms. Isoform II initiates at TSS2 and encodes a 380 aa protein with a predicted mitrochondrial targeting sequence (MTS; Mitoprot score = 0.9971; (Claros and Vincens, 1996). Isoform I initiates at TSS1 and after splicing encodes a 337 aa protein lacking the MTS. Right: Primer extension (primer location indicated by red splotch) reveals that isoform II predominates during log phase growth, but isoform I appears upon stress [cycloheximide (CHX) or late log/glucose deprivation]. See also Figure S2 and Table S1 and S2

### Correcting intron annotations, alternative 5’ SS’ s and recursive splicing

In addition to the 5’-P and 3’-OH peaks discussed above, many other peaks met the <0.01 FDR threshold but did not correspond to current splice site annotations. Direct examination of all such peaks with mean >50 reads per million mapped (rpm; 328 peaks total) revealed numerous misannotated or previously unannotated spliced sites that we were able to confirm by parallel examination of exon junction reads in existing WT RNA-Seq data (Marguerat et al., 2012; Rhind et al., 2011) and/or spliceosome RNA-Seq and footprints. In all we found sixteen 5’SS and 3’SS misannotations in CDS genes, all of which change the encoded protein (**Table S2A**). Of these sixteen, nine result in a small amino acid insertion or deletion (1-28 aa) at an internal position, whereas five were due to transcription start site (TSS) or intron misannotations at the end of a CDS region (where genome annotation is inherently more difficult).

Two particularly interesting examples of previously misannotated introns within CDS regions occur in transcripts encoding spliceosomal proteins. One is an intron initiating with A and terminating with C (aka, an AT-AC intron) in SPBC1861.08c (Lea1) (**Figure 2B**). Although *S. pombe* does not contain the U11/U12-dependent minor spliceosome responsible for removing most AT-AC introns in metazoans, the U1/U2-dependent major spliceosome can remove introns starting with A and ending with C in the context of otherwise strong 5’SS, BP and 3’SS consensus sequences (Wu and Krainer, 1997). We confirmed this new AT-AC annotation by Sanger sequencing of the RT-PCR cDNA product generated from untagged WT *S. pombe* RNA (**Figure 2B inset**) and examination of WT RNA-Seq reads (**Figure 2B bottom**). Another interesting example was a 3’SS misannotation in SPAC27F1.09c (prp10/sap155/SF3B1) intron 2 (**Figure 2C**). SF3B1 is both an essential spliceosomal protein and an oncogene commonly mutated in cancers (Jenkins and Kielkopf, 2017). Untagged WT cells express two prp10 mRNA isoforms, one retaining the intron and encoding the full-length protein and another where the corrected intron has been removed and the CDS frameshifted (**Figure 2C inset**). The abundance of the frameshifted isoform was higher in cells lacking Upf1, an essential nonsense-mediated decay (NMD) factor. Thus it appears that, like metazoans (Ge and Porse, 2014; Jacob and Smith, 2017), *S. pombe* uses alternative splicing linked to NMD (AS-NMD) to maintain homeostasis of this crucial splicing factor.

Finally, unexpected peaks in the 336 nt intron in SPNCRNA.445 (snoRNA61) revealed an internal zero-length exon (i.e., a 3’SS immediately adjacent to a 5’SS) (**Figure 2D**). RT-PCR confirmed complete removal of the full-length intron. In multicellular organisms, zero-length exons are sites at which recursive splicing enables intron removal piecewise (Georgomanolis et al., 2016). To our knowledge, these are the first documented example of recursive splicing in a yeast species. Sequence conservation (Rhind et al., 2011) in other *Schizosaccharomyces* species of SPNCRNA.445 intron 1 including the splice site sequences surrounding the zero-length exons, suggests that recursive splicing of these introns serves an important biological purpose.

### New introns

Our data also revealed 217 new introns. Eighteen of these occur in previously unannotated transcripts (**Table S2B**) and 103 interrupt ncRNAs of unknown function (**Table S2C**). Sixteen others harbor snoRNAs, including five new introns in the SPNCRNA659 host gene, each surrounding snoRNAs 2, 3, 4, 5 and 6 (**Figure S2**). Whereas the majority (62 of 80) of novel introns in CDS genes lie fully within UTRs (**Table S2D**), nineteen reside either partially or fully within the annotated ORF and so alter the predicted protein sequence. One particularly interesting example is a new 543 nt intron overlapping the 5’-UTR and ORF of Alcohol dehydrogenase 4, adh4 (SPAC5H10.06c) (**Figure 2E**). Primer extension and RT-PCR analyses indicated that adh4 has two mRNA isoforms produced from alternative transcription start sites (TSS1 and TSS2), with TSS2 residing within the newly detected intron. PolyA+ RNA-Seq tags indicative of both mRNA isoforms were readily identifiable in previous datasets from untagged WT cells (Marguerat et al., 2012). *S. pombe* Adh4 protein was previously shown to co-purify with mitochondria (Crichton et al., 2007), and transcripts initiating at TSS2 encode a protein isoform (II) containing an N-terminal mitochondrial targeting sequence (MTS). Initiation at TSS1, however, leads to intron excision and removal of the first 43 codons containing the MTS – thus protein isoform I is presumably cytoplasmic. The relative abundance of these two mRNA isoforms depends on growth conditions (**Figure 2E**). During log phase growth in rich media, mRNA isoform II predominates, but upon stress (e.g., inhibition of translation or glucose deprivation), mRNA isoform I predominates. Thus, in addition to identifying a new intron in adh4, our 5’-P and 3’-OH sequencing led us to discover a stress-responsive TSS and splicing switch that likely alters the subcellular targeting of a key metabolic enzyme.

### Retained and high splicing intermediate introns

For prp10, the WT RNA-Seq data and our RT-PCR analysis indicated a significant cDNA fraction retaining the queried intron (**Figure 2B, C**). This led us to examine the relationship between different intron populations and their retention levels in RNA-Seq libraries. Introns covered by our 5’-P or 3’-OH datasets and for which the ASM294v2.26 genome annotation agreed with our spliceosome profiling data (i.e., the vast majority of all previously annotated introns) tended to have very low intron retention percentages in untagged WT RNA-Seq datasets (**Figure 3A, B**). Newly detected introns, however, tended toward higher intron retention, likely explaining why they were previously missed in annotations based on RNA-Seq alone.

**Figure 3.**
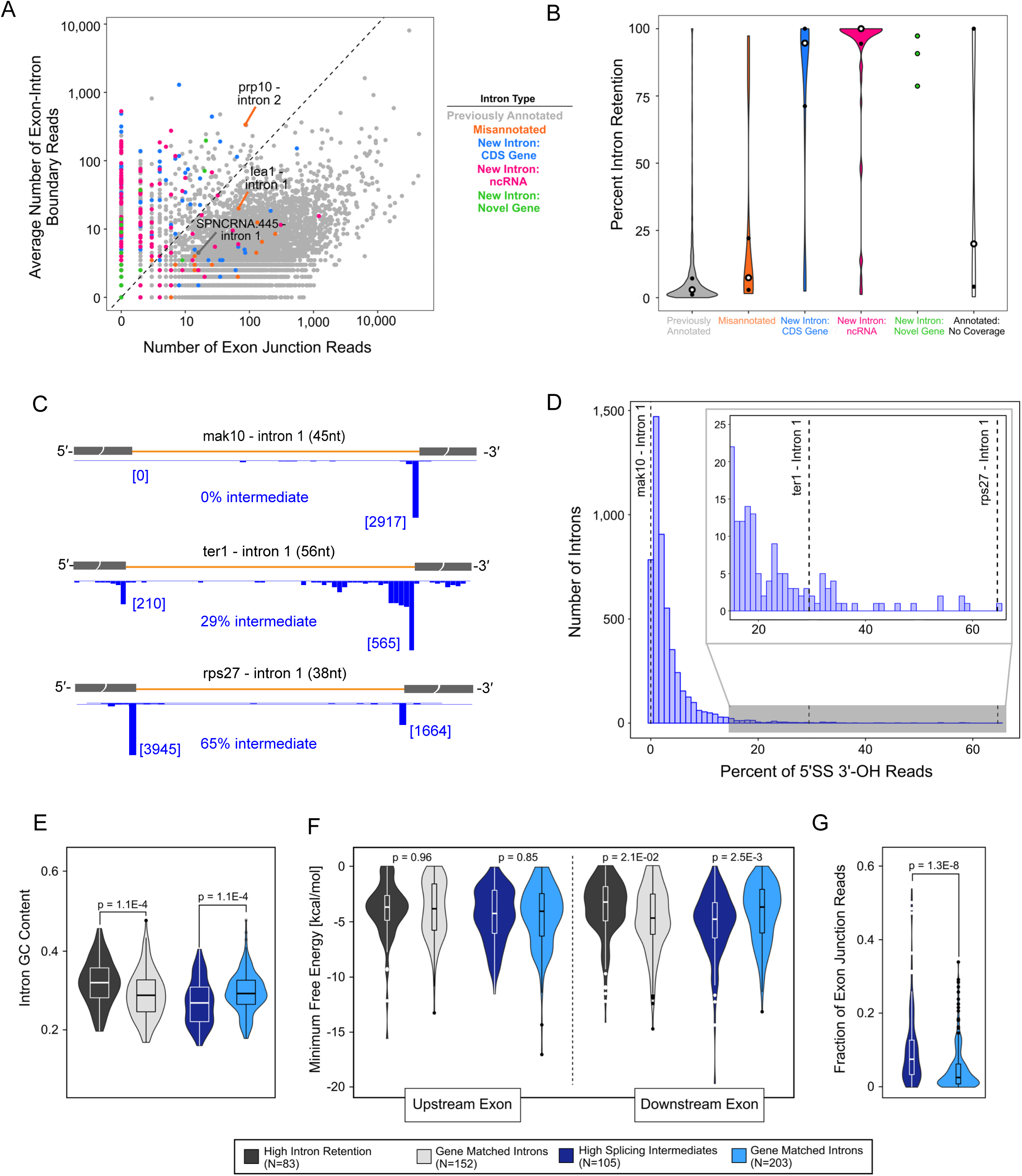
Retained and high splicing intermediate introns. (A) Scatterplot of the number of exon junction reads verses number of exon-intron boundary reads from WT RNA-Seq data for all *S. pombe* introns colored by type as indicated. Introns previously mentioned in Figure 2 are labeled. (B) Violin plots comparing percent of intron retention between intron groups defined in A. Previously annotated introns not covered by 5’-P or 3’-OH data are shown in white. (C) Examples of introns with different levels of splicing intermediates. Numbers in brackets: 3’-OH 3’ read ends terminating at the 5’SS or 3’SS. (D) Histogram of percent of splicing intermediate (calculated as fraction of 5’SS 3’-OH to all intron 3’-OH reads) across all introns with ≥ 25 3’SS 3’-OH reads in both replicates. Dashed vertical lines indicate introns shown in A. (E) Violin plots comparing GC content of introns with high intron retention percentages (>70%) to all other introns in the same gene (gene matched introns) and introns with high (>17%) splicing intermediate to gene matched introns. P-values: Wilcoxon rank sum test. (F) Violin plots comparing folding free energy (RNAFold) for upstream (−1 to −40 nts from 5’SS) and downstream (+1 to +40 nts from 3’SS) exon sequences for sets described in E. P-values: Wilcoxon rank sum test. (G) Violin plots of fraction of exon junction reads for introns with high splicing intermediate to gene matched introns. P-values: Wilcoxon rank sum test. See also Figure S3 and Table S1 and S3

Introns can vary with respect to either first step efficiency (leading to varying degrees of full intron retention) or second step efficiency (leading to varying splicing intermediate abundance). In addition to the major 3’-OH peak at the 3’SS (3’SS 3’-OH reads) indicative of a fully excised intron, 3,528 introns had a second smaller 3’-OH peak (binomial test, FDR < 0.01, **Table S3A**) at the 3’ end of the upstream exon (5’SS 3’-OH reads) indicative of splicing intermediate (**Figure 2A**). Across the 5,110 introns (old plus new annotations) having at least 25 3’SS 3’-OH reads in both biological replicates, 5’SS 3’-OH reads averaged 3.3% of total (3’SS + 5’SS) 3’-OH reads for that intron, indicating that endogenous U2·U5·U6·NTC complexes contain ∼3% splicing intermediates on average, validating our previous estimate (Chen et al., 2014). Nonetheless, there was much variability among introns. For example, whereas the sole intron in mak10 (SPBC1861.03) had a total of 2,717 3’SS 3’-OH reads in the two biological replicates combined, no 5’SS 3’-OH reads were detected. At the opposite extreme, 5’SS 3’-OH reads were ∼2-fold more abundant than 3’SS 3’-OH reads for rps27 (SPBC1685.10) intron 1, indicating the presence of 65% splicing intermediate (**Figure 3C**).

High levels of splicing intermediate can indicate inefficient exon ligation, the means by which the 3’-end of telomerase RNA is generated from the *S. pombe* ter1 transcript (Box et al., 2008). Consistent with this, we observed 29% splicing intermediate on the ter1 intron (**Figure 3C**), ranking it within the top 2% of all 5,110 introns with regard to percent splicing intermediate (**Figure 3D**). Other ncRNA transcripts included in the top 100 introns exhibiting high (>17%) splicing intermediate included SPNCRNA.445 (snoRNA 61), SPSNORNA.01 (snoRNA 40), SPSNORNA.30 (snoRNA 62), and three transcripts antisense to CDS genes (**Table S3A**). In the future it will be of interest to determine whether any of these generate functional RNA species through inefficient exon ligation.

All other high splicing intermediate introns were in CDS genes. Gene ontology analysis of the top 100, 150 or 200 introns ranked by fraction of splicing intermediate revealed them to disproportionately interrupt genes encoding ribosomal proteins (RPs) (**Table S3B-D**). Interestingly, this gene ontology enrichment did not extend to CDS genes with high levels of full intron retention (**Figure 3A**). There was also very little overlap between the high intron retention and high splicing intermediate CDS gene sets (**Figure S3A**), and no specific gene ontology terms were associated with the high intron retention set. In budding yeast, inefficient intron excision from RP transcripts serves to negatively regulate ribosome production (Juneau et al., 2006; Parenteau et al., 2011). Our findings thus suggest that this may also be the case in *S. pombe*, with the regulation occurring at the second step of splicing, not initial intron recognition.

Examination of spliceosome footprinting reads across introns with high splicing intermediate revealed that many also had a high fraction of exon junction reads (**Figure S3B, C**). A high fraction of spliced exons could indicate inefficient spliced product release. Because the spliceosome can catalyze both forward and reverse splicing (Tseng and Cheng, 2008), inefficient spliced exon release could result in reverse second step, and therefore an increase in splicing intermediates. Consistent with this idea, introns with the highest fraction of splicing intermediates had statistically higher fractions of exon junction reads than all other introns within the same genes (gene-matched intron set) (**Figure 3C**).

To identify intron and exon features associated with high intron retention, high splicing intermediate and/or high spliced exon junction reads, we compared the 100, 150 and 200 top intron sets defined above against similarly sized sets of (1) introns with the lowest percent of intron retention or splicing intermediates, respectively, or (2) a gene-matched intron set (**Figure 3E-G, Figure S3**). Features analyzed included 5’SS, BP and 3’SS motifs, intron length, intron order, BP to 3’SS distance, sequence conservation among *Schizosaccharomyces* species, essential vs nonessential genes, intron and exon nucleotide content and intron and exon folding energies. For highly retained introns, the only statistically significant differences were higher intron GC content (p<10^−4^) and less predicted structure in the downstream exon (p<0.02). In metazoans, introns tend to have equal or lower GC content than the flanking exons (Amit et al., 2012). Thus high intron GC content may interfere with initial intron recognition. Conversely, high splicing intermediate introns had lower GC (p<3×10^−4^) and higher A (p<10^−8^) content within the intron, higher GC-content in the flanking exons (p<0.004), and a stronger predicted folding energy (Lorenz et al., 2011) of the downstream exon (p<0.002). We also note that strong secondary structure driven by high GC-content in a downstream exon could inhibit binding of Prp22, the DExH-box RNA helicase responsible for releasing the spliced exons (Schwer, 2008).

### Testing the branch-only model for 5’ SS cleavage

The currently accepted view of spliceosome catalysis is that 5’SS cleavage occurs solely by attack of the BP adenosine 2’-OH on the 5’SS phosphodiester to liberate the 5’ exon and generate a lariat intron containing a 2’-5’ bond between the BP and 5’SS. However, group II self-splicing introns, which are both evolutionarily and structurally related to the spliceosome (Pyle, 2016), can catalyze 5’SS cleavage by either branch formation or 5’SS hydrolysis, the latter resulting in a linear intron-3’ exon intermediate (Jarrell et al., 1988; Podar et al., 1998). As discussed in the Introduction, the majority of *S. pombe* and metazoan introns still lack a positively identified BP. A potential explanation was that some spliceosomal 5’SS cleavage events occur by hydrolysis rather than branching.

To investigate the possibility of 5’SS hydrolysis, we made 5’-P and 3’-OH libraries as above, but without DBE treatment (DBE-). About 34% of introns (1,907) behaved as expected, with a 5’SS 5’-P peak only detectable after DBE treatment (**Figure 4A** and **Figure S4A**). The other 66% (3,686), however, had a significant (albeit reduced) 5’SS 5’-P peak in at least one DBE-replicate (**Figure 4B** and **Figure S4A**). To investigate the possibility that some introns are subject to debranching by endogenous debranching enzyme (Dbr1) prior to U2·U5·U6·NTC complex disassembly, we isolated U2·U5·U6·NTC complex from dbr1Δ cells and prepared DBE+ and DBE-5’-P and 3-OH libraries. About 8% of introns still retained significant 5’SS 5’-P peaks across both biological replicates even in the absence of both endogenous DBR1 and exogenous DBE (**Figure 4B** and **Figure S4B**). Incubation of ^32^P-labeled branched oligos in cell extracts confirmed that, while debranching activity was clearly apparent in the parental TT extract, the dbr1Δ extract had no detectable debranching activity (**Figure 4C**). Thus a fraction of introns co-purifying with U2·U5·U6·NTC complex are linear and these linear species are not produced by Dbr1.

**Figure 4.**
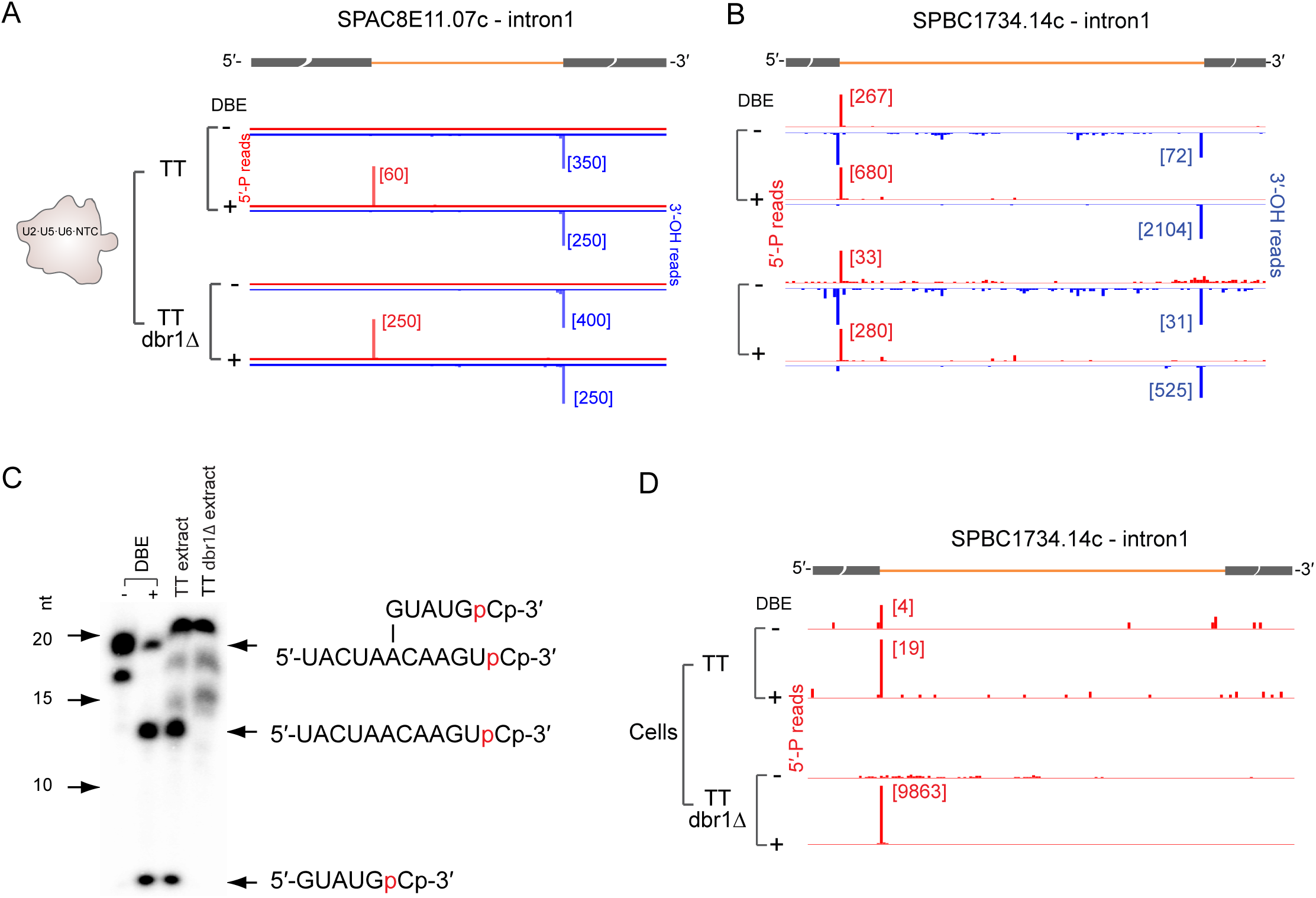
Dbr 1 - independent debranching in purified U2·U5·U6·NTC complexes. (A) 5’-P and 3’-OH read ends on SPACBE11.07c intron 1 copurifying with U2·U5·U6·NTC complexes from TT or TT dbr1⊗ cells with (+) or without (−) DBE treatment. Observation of a 5’SS 5’P peak only upon DBE treatment indicates a high proportion of lariat species. (B) Same as A, except SPBC1734.14c intron 1. Observation of 5’-P peak in all samples indicates that a fraction of intron-containing species co-purifying with U2·U5·U6·NTC complexes are linear and are not due to Dbr1 activity. (C) Polyacrylamide gel showing migration of 3’-end labeled branched oligo (structure at right) and cleavage products without (−) or following DBE treatment (+) or incubation in TT or TT dbr1⊗ cell extract. Lack of cleavage products in the dbr1⊗ cell extract indicates absence of another cellular debranching activity. (D) Same as B, except 5’-P reads from cellular RNA. See also Figure S4 and Table S1

Under certain salt conditions, purified human spliceosomes are capable of hydrolyzing the 2’-5’ bond at the BP (Tseng and Cheng, 2013). Thus the possibility remained that the linear species in the dbr1Δ U2·U5·U6·NTC complex were generated during our spliceosome purification protocol. To test this, we made DBE+ and DBE-5’-P libraries from total cellular RNA extracted from both the TT and dbr1Δ strains. In the DBE+ libraries, 4,705 introns had statistically significant (FDR < 0.01) 5’SS 5’-P peaks in both TT and dbr1Δ, and 3,640 5’SS 5’-P peaks were detectable in DBE-TT cellular RNA. Notably, however, no statistically significant 5’SS peaks were found in the DBE-dbr1Δ cellular RNA (**Figure 4D**). Thus the linear introns present in purified dbr1Δ U2·U5·U6·NTC complexes were most likely due to 2’-5’ bond hydrolysis occurring during spliceosome purification and not initial 5’SS cleavage by hydrolysis. We conclude that, unlike group II introns which can activate either H_2_O or the BP 2’-OH as the nucleophile for the first step of splicing, the spliceosome can only initiate 5’SS cleavage by activating the BP 2’-OH. Therefore the inability to positively identify BPs in many introns is most likely due to the inherent inefficiencies in previous transcriptome-wide mapping methodologies.

### Spliceosome footprints

Recent cryo-EM structures of spliceosomes have yielded unprecedented views into how the snRNAs and core spliceosomal proteins are arranged in 3-dimensions (Shi, 2017). The longest intron sequences modeled into the structures to date include the first 15 nts downstream of the 5’SS and 23 nts around the BP (14nts-UACUAAC-2nts) (Yan et al., 2016). To investigate whether this represents the full extent of strong intron-spliceosome interactions, we performed footprinting experiments akin to ribosome footprinting. Briefly, during the U2·U5·U6·NTC complex purification protocol, we included a micrococcal nuclease (MNase) digestion step; we then treated the purified RNAs with DBE and made footprint libraries using a recently published lab protocol (Heyer et al., 2015) (**Figure 1B**). Single end sequencing (82 nts) of two biological replicates yielded 20 to 32 million reads (**Table S1A**) with captured fragment length ranging from 6 to 67 nts. As expected, a high percentage of reads mapped to U2, U5 and U6 snRNAs (∼70%), but of the remaining 1.5 million reads that mapped uniquely to the genome, 82% mapped to introns and exon-exon junctions.

Examination of reads on individual genes consistently revealed one peak on shorter (<60 nts) introns, a broad peak on intermediate length (∼60-150 nts) introns and two or three peaks on longer (>150 nts) introns (**Figure 5A**). To analyze these footprints in greater detail, we binned according to length all 4,746 annotated introns that were 35-200 nts long (excluding snRNA introns). Roughly half (2,627) are ≤56 nts long; for these, there is a near perfect correlation (*r*=0.96) between the average number of footprint reads per intron and intron length (**Figure 5B**). For introns longer than 56 nts, the average number of footprint reads per intron was largely independent of intron length. This suggests that our purified splicing complexes protect a total of 56 intronic nts. For introns ≤56 nts, this protected region encompasses the entire intron.

**Figure 5.**
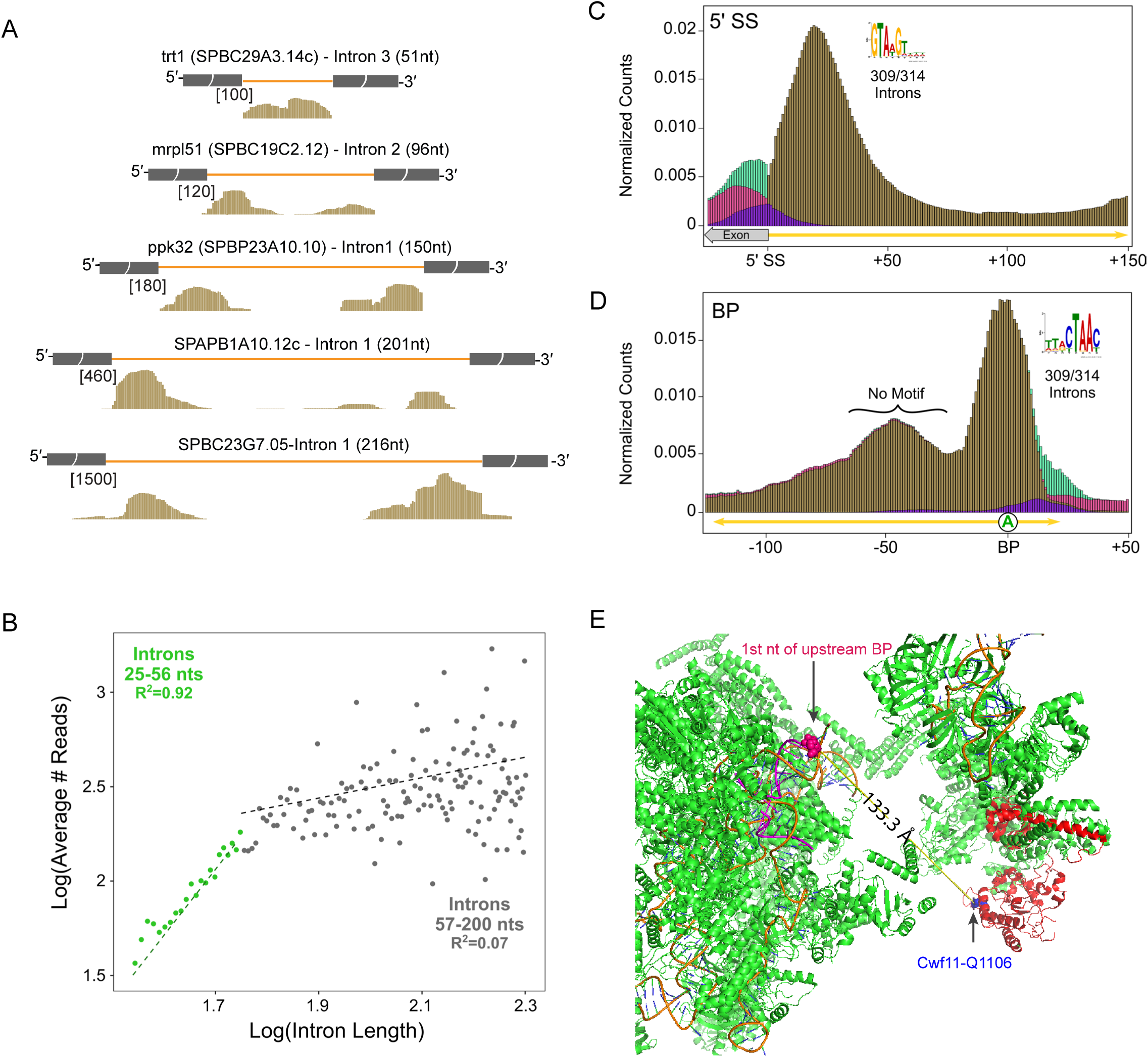
Spliceosome footprints. (A) Examples of spliceosome footprints on short (51 nt), medium (96 and 150 nt), and long (210 and 216 nt) introns. Intron identities are as indicated, with scales in brackets. (B) Scatter plot showing average footprint read counts for all introns in each one nt length bin. (C) Aggregation plot of normalized read counts (number of reads at position divided by total number of intron reads from both footprinting library replicates) mapping to the end of the 5’ exon and/or within the first 150 nts for 314 introns with length >200 nts. Brown: Counts from reads mapping fully within the intron; Purple: Counts from reads spanning the 5’SS; Pink: Counts from reads mapping fully within the exon; Green: Counts from exon junction spanning reads. Inset: sequence logo for 5’SS motif found *de novo* using MEME. (D) Same as D, except numbers indicate nucleotide positions relative to BP identified using FIMO and consensus motif. Whereas MEME identified a strong motif over the BP, no motifs were detectable in the footprint centered 48-49 nts upstream. (E) Partial structure of the *S. pombe* U2·U5·U6·NTC complex (Yan et al., 2015) showing distance between BP and RNA binding motif in Cwf11. See also Table S1

To cleanly separate the 5’SS and 3’SS footprints and determine their lengths, we made aggregation plots of reads mapping to the 5’ and 3’ ends of introns that were >200 nts (**Figure 5C,D**). At both ends, the peaks were rather broad, indicating that the length of RNA protected by the spliceosome is somewhat variable. The peak summit downstream of the 5’SS occurs at +19 nts and is half maximal between +5 and +36 nts (**Figure 5C**). At intron 3’ ends, footprints center on the TACTAAC branch site consensus, extending 14 nts upstream and 11 nts downstream for a total of 25 nts; in many introns, this protected region extends all the way to the 3’SS (**Figure 5D**). Thus, the spliceosome protects ∼31 nts at the 5’ ends of introns and ∼ 25 nts around the BP.

On long introns, we also observed a third peak centered 48-49 nts upstream of the BP with a breadth of ∼38 nts (**Figure 5A, D**). *De novo* motif analysis using the 314 introns in this set revealed no specific motifs; thus its presence is not due to alternative 5’SSs or BPs. More likely this peak represents the binding site of Cwf11 (aka IBP160, Aquarius), an SF1 RNA helicase known to interact in a sequence-independent manner with the region upstream of the BP (**Figure 5E**).

### BP identification: B reads and BAL algorithm

Although intron ends are readily defined from RNA-Seq data, BP locations are more challenging. If reverse transcriptase initiates cDNA synthesis near the 5’ end of a lariat intron, it can cross the BP and generate a read wherein the region upstream of the BP is concatenated with the region downstream of the 5’SS to generate a branched (“B”) read (**Figure 6B**). These B reads often contain a mismatch or indel at the BP nucleotide due to the presence of a 2’-5’ bond (Gao et al., 2008; Vogel and Borner, 2002; Vogel et al., 1997). Since our purified spliceosomes should be highly enriched for BPs, we developed a <u>B</u>ranch <u>AL</u>igner (BAL) algorithm similar to one previously described for mapping human BPs (Mercer et al., 2015) and applied it to spliceosome RNA-Seq and footprinting libraries constructed from RNAs copurifying with U2·U5·U6·NTC complexes from both TT and dbr1Δ strains (**Figure 6A**). In all, we identified 4,389 B reads from a total of 186,863,605 reads, enabling us to map 1,463 BPs in 1,420 introns. The vast majority (98.6%) of these B reads had a mismatch at the BP. While many of BAL-defined BPs overlapped with those previously identified in studies also employing B reads (Awan et al., 2013) (Bitton et al., 2014) (**Figure 6C**), 770 were new. All three studies combined resulted in positive identification of 3,759 BPs in 2,374 of *S. pombe’s* 5,533 spliceosomal introns (∼43% of total). However, the high fraction of non-overlapping BPs among the three studies indicated that none achieved saturation.

**Figure 6.**
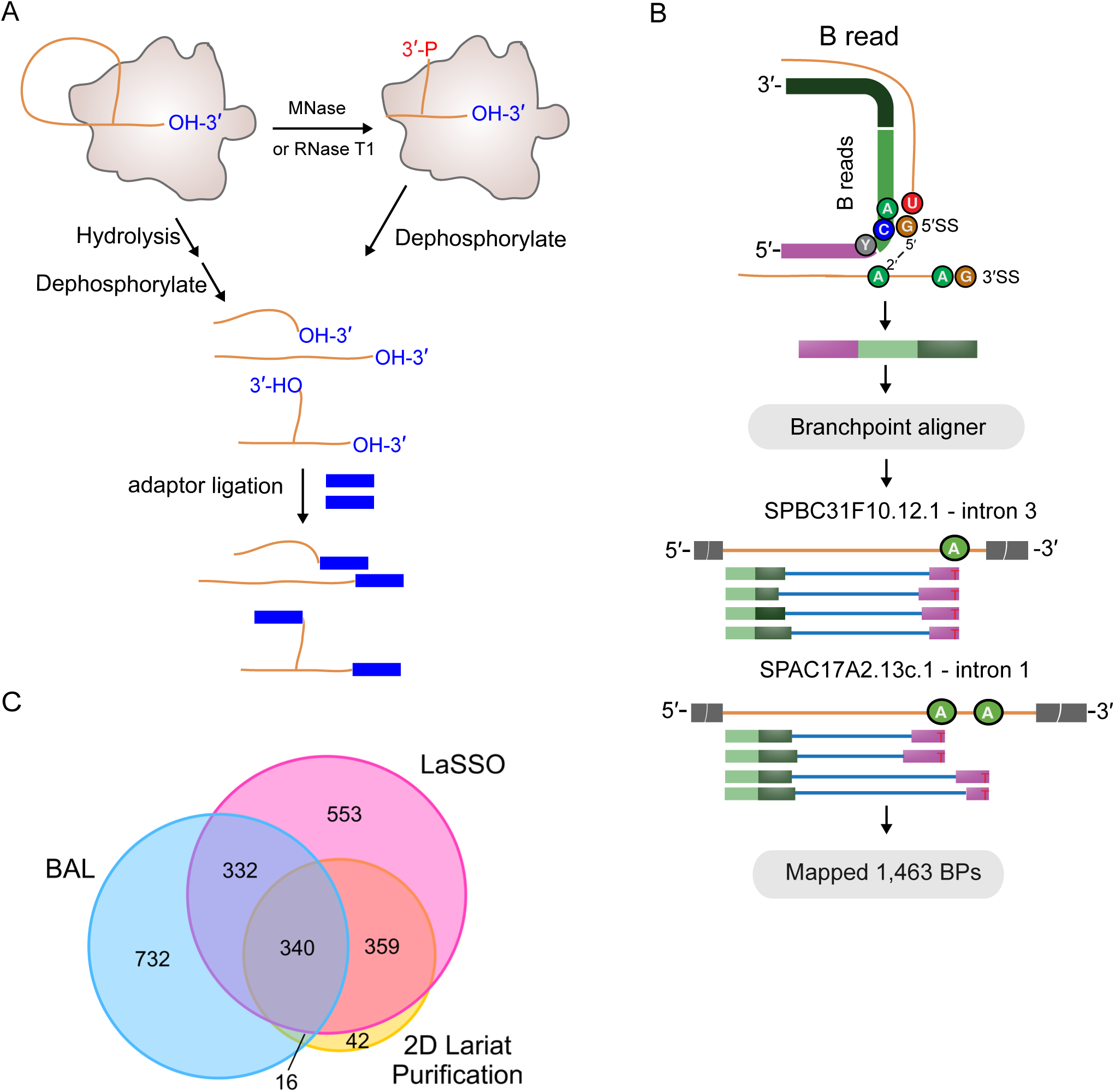
BP identification by B reads and BAL algorithm. (A) Schematic of libraries used to identify B reads. (B) Top: Schematic of BAL pipeline showing 20 nt seed (light green) downstream of 5’SS, seed extension (dark green) and 5’ remainder mapping upstream of the BP (purple). Bottom: Examples of BPs supported by B reads. (C) Overlap of introns with BPs identified by BAL (blue), LaSSO (pink; (Bitton et al., 2014)), and 2D gel lariat purification (yellow;(Awan et al., 2013)). See also Table S1 and S4

### BP identification: G reads and random forest approach

As the above numbers illustrate, a major limitation to detecting branches via B reads is their extreme scarcity, even in libraries highly enriched for introns (one per 200,000 mapped reads in our datasets). This is likely due to the propensity of SuperScript III reverse transcriptase to stop at a BP rather than cross the 2’-5’ bond and continue on. In our libraries the ratio of B reads to reads ending at the 5’SS was 0.0075, meaning that SuperScript III stopped >99% of the time when it encountered a 2’-5’ bond. Much more common in our DBE-datasets were “G reads”: <u>g</u>enome-mapping reads either crossing or ending at the BP (**Figure 7A** and **B**). In the dbr1Δ spliceosome RNA-Seq and 3’-OH libraries 8% of all genome-mapping reads (2,202,687 of 26,669,756 total) ended at a 3’SS (i.e., came from the intron product). For the introns with a BP identified by both BAL and Anwar *et al*. (2012), 15% of these crossed the BP, 44% began at the BP nucleotide and 32% began one nucleotide downstream of the BP. The remaining 9% began between the BP and 3’SS, so were uninformative as to BP location. Thus, SuperScript III is substantially more efficient at crossing the 3’-5’ bond at the BP (15% read through) than it is at crossing the 2’-5’ bond (0.075% read through), and 7.3% (= 8% × 91%) of all genome-mapping reads in the dbr1Δ spliceosome RNA-Seq and 3’-OH libraries contained information as to a BP location.

**Figure 7.**
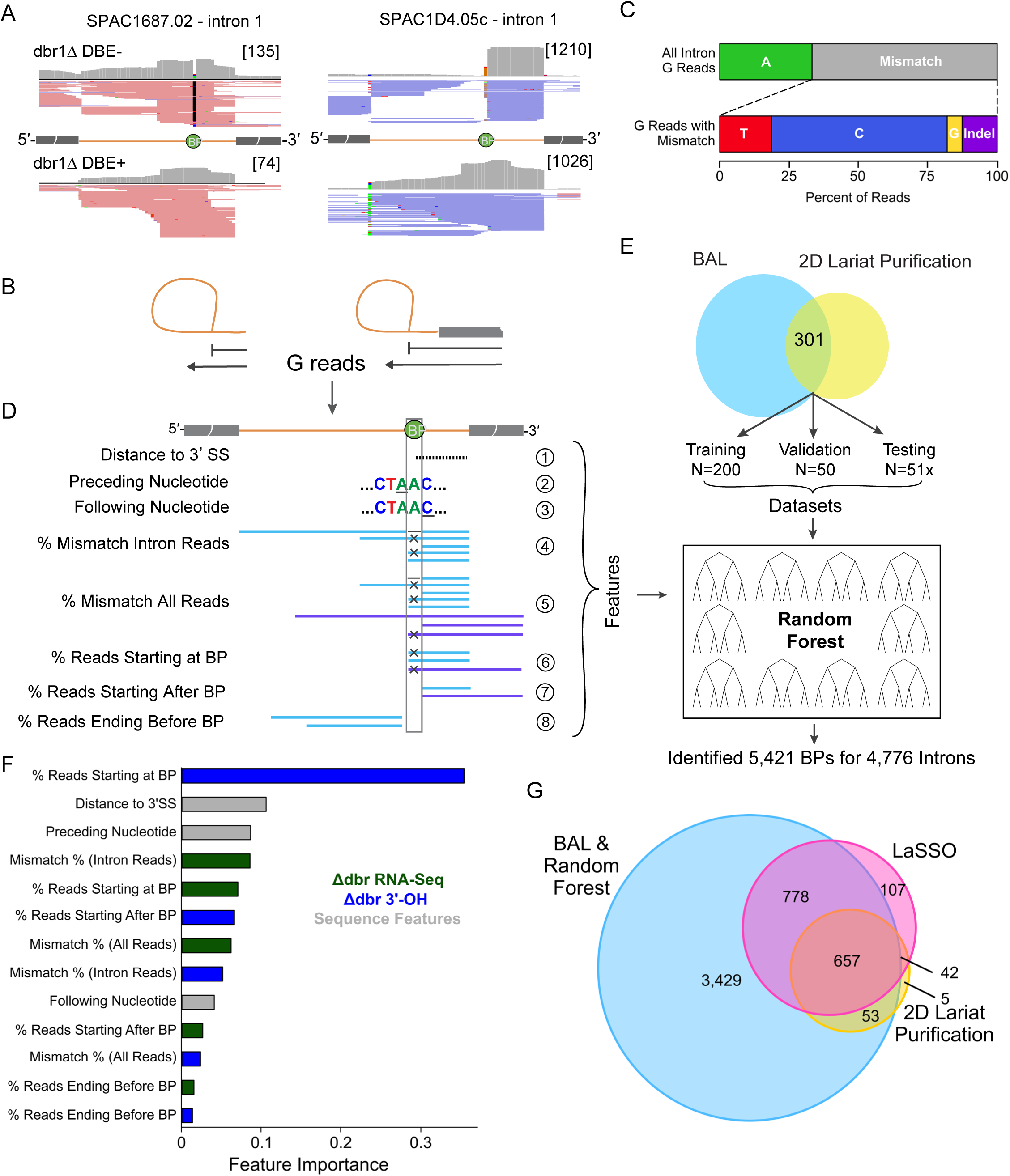
BP identification by G reads and random forest approach. (A) Examples of spliceosome RNA-Seq reads used to identify BPs. Upper left: When reads in DBE-libraries traverse the BP, they often contain a mutation at BP (black). Upper right: Other DBE-reads terminate at or one nucleotide before the BP. Lower: These patterns are not observed in DBE+ libraries. (B) Schematic of genome mapping “G” reads that can be used to identify BPs. Such reads can derive from lariat product (left) or lariat intermediate (right). (C) Mismatch rates of G reads crossing previously detected BPs (i.e., identified by both BAL and 2D gel lariat purification). (D) Features used in Random Forest Model. (E) Overview of Random Forest Model. Using the features in D, we trained the model on 200 BPs identified by both BAL and 2D gel lariat purification. We tuned parameters using a 50 BP validation set and evaluated model performance with a 51 BP test set. (F) Importance of features from Random Forest Model. Features are colored based on whether they are derived from the spliceosome RNA-Seq data (green), 3’-OH data (blue), or from the sequence itself (gray). (G) Overlap of the number of introns with BPs identified by BAL or Random Forest Model (blue), LaSSO (pink), and 2D gel lariat purification (yellow). See also Figure S5 and Table S1 and S4

As with the B reads, we found that SuperScript III has a strong tendency to make a mistake (point mutation or indel 67% of the time) when traversing the 3’-5’ bond at the BP (**Figure 7C**). It therefore seemed likely that we could use this information along with other features (e.g., BP sequence motifs, distance to 3’SS) to identify additional BP’s via a supervised machine learning approach (**Figure 7D**). Using the combined reads from all U2·U5·U6·NTC complex dbr1Δ DBE-spliceosome RNA-Seq and 3’-OH datasets (4 in total) and a set of 251 highly validated branch points (i.e., identified by both BAL and (Awan et al., 2013)), we generated and validated a model using the Random Forest algorithm (Breiman, 2001) (**Figure 7E**). When challenged with a 50 BP test set, this model exhibited remarkably high performance (AUROC = 0.99; AUPR= 0.98; **Figure S5A**), outperforming a BP identification method using solely the BP consensus motif (AUROC= 0.98, AUPR= 0.82; **Figure S5B**). Among all features in our model, the most important was the fraction of 3’-OH reads terminating at the BP, followed by distance from the BP to the 3’SS (**Figure 7F**).

Using our Random Forest model, we were able to identify 5,421 BPs in 4,776 introns (86%) (**Figure S5C**). The previous branch point datasets (Pleiss and LaSSO) are largely included in our set (**Figure 7G**), with the combined set representing 5,045 of 5,533 total introns (91%) (**Table S4**). In most aspects (e.g., BP motif, distance to 3’SS) BPs identified by our Random Forest approach were indistinguishable from previously identified BPs (**Figure S5DE**). However, whereas all B read-only methods favored BP identification in longer introns, our Random Forest approach showed no such intron length bias (**Figure S5F**). Thus, taking advantage of the G reads contained in deep sequencing libraries prepared from purified, lariat-containing spliceosomes allows for much more comprehensive BP identification transcriptome-wide than approaches based on B reads alone.

## Discussion

Recent implementation of “ribosome profiling” has greatly advanced the study of general and regulated translation. That technique, which involves deep sequencing of the mRNA footprints underlying translating ribosomes, is transforming our understanding of post-transcriptional gene regulation in the cytoplasm (Brar and Weissman, 2015; Ingolia, 2014). By analogy, we hereby dub the deep sequencing of RNAs co-purifying with spliceosomes as “spliceosome profiling”. Because it provides a means to directly interrogate all sites of spliceosome assembly and catalysis genome-wide, we expect that spliceosome profiling will similarly transform our understanding of post-transcriptional gene regulation in the nucleus.

Depending on the type of deep sequencing library constructed, spliceosome profiling can provide many different types of information. For example, spliceosome RNA-Seq libraries (i.e., containing long random fragments of introns) enable visualization of entire introns (**Figure 1**). Exactly where on the introns spliceosomes reside can be revealed by including a nuclease digestion step to generate spliceosome footprints (**Figure 5**). Creation of 5’-P and 3’-OH libraries after debranching allows for the unambiguous identification of intron ends at nucleotide resolution, discovery of previously unannotated introns and alternative splice sites (**Figure 2**), and relative quantitation of splicing intermediates versus products (**Figure 3**). Finally, a new random forest approach for detecting BPs applied to various libraries from samples not subjected to debranching allows for intronome-wide BP identification at unprecedented depths (**Figure 7**).

In the work reported here, our analysis focused on RNA species co-purifying with a Protein A-tagged U2 component and a CBP-tagged U5 component in *S. pombe* during late log growth in rich media. This tandem affinity approach purifies endogenous U2·U5·U6·NTC complexes, which mainly consist of spliceosomes containing fully excised lariat introns (ILS complex). Profiling of these complexes thus provides a transcriptome-wide snapshot for the very last stage of spliceosome action. However, many additional spliceosome assembly states exist in cells, including cross-exon complexes and cross-intron U1·U2 Complex A (aka, pre-spliceosomes), U1·U2·U4·U5·U6 pre-activation Complex B, U2·U5·U6·NTC post-activation Complex B^act^, U2·U5·U6·NTC splicing intermediate-containing Complex C, and U2·U5·U6·NTC splice product-containing Complex P. Extensive proteomics analyses of these different complexes assembled in vitro have revealed that each has a unique protein signature (Fabrizio et al., 2009; Wahl et al., 2009). Therefore, by judicious choice of protein pairs it should be possible to query nearly every spliceosome assembly state genome-wide by tandem affinity purification or RNA immunoprecipitation in tandem (RIPiT) (Singh et al., 2012; 2014). Gene editing advances now make it possible to insert affinity tags into almost any organism (Lackner et al., 2015). Thus, our overall approach should enable generation of genome-wide occupancy maps for all major spliceosome assembly states in diverse cell types. We predict that spliceosome profiling will prove especially valuable for elucidating the mechanisms governing alternative pre-mRNA processing among different tissues of multicellular organisms or in cells subject to different growth, signalling or stress conditions.

### New sites of spliceosome action

Our analysis via 5’-P-Seq and 3’-OH-Seq of intron ends copurifying with endogenous U2·U5·U6·NTC complexes enabled us to identify >200 new *S. pombe* introns and correct the annotation of numerous others (**Figure 2 and Table S2**). Many of the new or corrected introns were in UTRs or near the end of an ORF, where computational algorithms for predicting gene structure are least reliable. Importantly, many corrected intron annotations change the predicted protein sequence, which can have significant consequences for protein localization and/or function. For example, a new intron arising from an alternative TSS in the adh4 gene encodes a protein isoform lacking the N-terminal mitochondrial targeting sequence (MTS) present in the previously annotated coding region. We showed that relative abundance of the two mRNA isoforms depends on cellular growth and stress conditions. Thus, spliceosome profiling led us to discover a stress-responsive transcriptional and splicing switch that alters the subcellular targeting of a key metabolic enzyme. Our data also revealed two instances of recursive splicing via zero-length exons, an AT-AC intron, a regulatory intron retention event in the gene for a key spliceosomal protein, and scores of new intron-containing transcripts (**Table S2B**). While these latter transcripts were readily detectable in our purified U2·U5·U6·NTC complexes, their spliced exon products were not apparent in traditional RNA-Seq libraries, possibly due to their rapid decay by nonsense-mediated decay (NMD). These findings demonstrate the utility of spliceosome profiling both for improving genome annotation and for identifying all sites of spliceosome catalysis regardless of intragenic position, matches to expected splice site consensus sequences or subsequent spliced exon stability.

In addition to allowing for intron identification and more accurate splice site mapping, 3’-OH read ratios at exon and intron ends provides a new window for assessing second step splicing efficiencies. Existing transcriptome-wide methods either detect splicing intermediates physically associated with RNA Pol II (Mayer et al., 2015; Nojima et al., 2015) or monitor overall splicing kinetics (Barrass et al., 2015; Bhatt et al., 2012). By using 3’-OH-Seq to specifically query exon and intron ends in catalytically active spliceosomes, our approach allows quantitative analysis of both first and second step chemistries (**Figure 3**) regardless of whether these events occur co- or post-transcriptionally. Using this approach we found that many introns with high levels of splicing intermediate occur within ribosomal protein genes, consistent with previous findings that splicing efficiency regulates ribosomal protein expression in *S. cerevisiae* (Juneau et al., 2006; Parenteau et al., 2011).

### Extensive spliceosomal footprints

Recent cryo-EM structure determination of late stage spliceosomes has enabled visualization of the 5’SS and BP spatial organization within the catalytic core (Galej et al., 2016; Wan et al., 2016; Yan et al., 2015). To date the structures containing the most modeled intron nucleotides are the *S. cerevisiae* C and C* complexes (Wan et al., 2016; Yan et al., 2016). In all, these models contain 15 nts downstream of the 5’SS and 23 nts surrounding the BP for a total of 38 nts. Consistent with this, our analysis of protected RNA fragments co-purifying with endogenous *S. pombe* U2·U5·U6·NTC complexes revealed a ∼31 nt footprint downstream of the 5’SS (**Figure 5C**) and a ∼25 nt footprint surrounding the BP (**Figure 5D**). Because the 5’SS and BP consensus sequences (<u>GU</u>AGA and UACA<u>A</u>C, respectively) are much shorter than the footprints, spliceosomal contacts with adjacent regions are presumably driven by sugar-phosphate backbone interactions. These structural contacts likely strengthen the association between the splicing machinery and intron so as to prevent consensus sequence dissociation during the many structural rearrangements required for spliceosome assembly and activation.

In addition to the 5’SS and BP footprints, we detected a third ∼38 nt footprint on long introns centered ∼38-39 nts upstream of the BP (**Figure 5D**). In human C complex, up to 44 nts 5’ of the BP are protected from RNase digestion (Wolf et al., 2012), with nts −33 to −40 interacting with Aquarius (AQR; aka IBP160/KIAA0560 in humans) (Hirose et al., 2006). A stoichiometric component of our purified U2·U5·U6·NTC complexes (Chen et al., 2014), the *S. pombe* AQR homolog, Cwf11, is thus likely responsible for this third footprint. Consistent with the lack of any detectable sequence motifs under the third footprint, AQR is an SF1 RNA helicase containing a sequence-independent RNA binding site spanning ∼ 7 nts (De et al., 2015; Hirose et al., 2006). In the recent cryo-EM structures of late stage *S. pombe* and human spliceosomes (Bertram et al., 2017; Yan et al., 2015), this RNA binding site is perfectly positioned relative to the spliceosomal catalytic core (∼14 nm away) to account for the observed footprint (**Figure 5E**).

Given the positioning of the Cwf11 footprint ∼50 nts upstream of the 3’SS (**Figure 5F**), an unresolved question is where does Cwf11 bind on introns not long enough to accommodate it (e.g., on the vast majority of S. pombe introns that are <60 nts)? In these cases does it simply not bind, or does it interact with the 5’ exon upstream of the 5’SS? In human C complex, the last 27 nts of the 5’ exon are fully protected from nuclease digestion, with a previously unidentified ∼170 kDa protein crosslinking in the region between the exon junction complex (EJC) deposition site and the 5’SS (Jurica et al., 2002; Reichert et al., 2002; Wolf et al., 2012). Intriguingly, in mammalian spliceosomes the 160 kDa AQR serves as a recruitment site for EJC components prior to their deposition on the 5’ exon (Ideue et al., 2007). Thus the ∼170 kDa crosslinked protein is most likely AQR.

### New methods for BP mapping

Although the BP consensus motif has high information content in budding and fission yeast, such is not the case in most metazoans (Lim and Burge, 2001). Thus, human BPs cannot be accurately predicted from sequence information alone. However, recent realization that mutations in spliceosomal proteins involved in BP selection are major drivers of cancer (Agrawal et al., 2018) has greatly increased the need for tools to precisely map human BPs. Our work here demonstrates that deep sequencing of RNA contained in late stage spliceosomes can increase BP identification to unprecedented depths. We used two different methods. The first, BAL, relied on identification of concatenated, branched (B) reads in our spliceosome RNA-Seq and spliceosome footprinting libraries. Although spliceosome purification should greatly increase the prevalence of such reads, we found that even in the footprinting libraries, where all sequences not stably protected by the splicing machinery were eliminated, only 4,389 of 186,863,605 (frequency = 2×10^−5^) reads corresponded to a B read. In comparison, 369,746 reads in the DBE-libraries ended exactly at the 5’SS indicating that the read-through efficiency of reverse transcriptase across a 2’-5’ bond is 0.75%. Consistent with B reads being insufficiently abundant to saturate coverage, we observed only partial overlap between the set of BPs we identified with BAL and previously identified sets (**Figure 6C**).

A much more successful approach was to use genome-mapping (G) reads that crossed or ended at the BP from the 3’ to 5’ direction. In our 3’-OH DBE-libraries, such reads represented 8% of all mapped reads. Our data indicate that when Super Script III initiates downstream of the BP, its read-through efficiency across the BP 3’-5’ bond (15%) is substantially higher its read-through efficiency across a 2’-5’ bond (0.75%). Further, when the BP 3’-5’ bond is traversed, there is a very high rate of mutation (67% of reads contain a point mutation or indel). By incorporating these features into a random forest algorithm (**Figure 7**), we were able to positively identify BPs on >86% of all introns in *S. pombe*. Importantly, the majority of all previously identified *S. pombe* BPs were included in this set. Thus our random forest approach using G reads from RNAs copurifying with late stage spliceosomes is a powerful new approach for BP identification. We anticipate that high-depth BP identification will be another useful application of spliceosome profiling.

## SUPPLEMENTAL INFORMATION

Supplemental Information includes five figures and four tables and can be found with this article online at…

## ACKNOWLEDGMENTS

We thank Beate Schwer and Masad Damha for their generous gifts of DBE and bRNA, respectively. We also thank Harleen Saini and Carrie Kovalak for critical reading of the manuscript. This work was supported by funding from HHMI, NIH R01-GM53007 (M.J.M.) and NSF DBI-0850008 (Z.W.). M.J.M. was a HHMI Investigator at the time this study was conducted.

## AUTHOR CONTRIBUTIONS

M.J.M. conceived and designed the experiments. W.C. performed the biochemical experiments. J.M., H.O. and H.P.S. performed all computational analyses. W.C., J.M., H.O., N.R., Z.W., and M.J.M. interpreted the data. W.C., J.M., and M.J.M. wrote the manuscript. All authors discussed the results and commented on the manuscript.

## Supplemental figure legends

**Figure S1.**
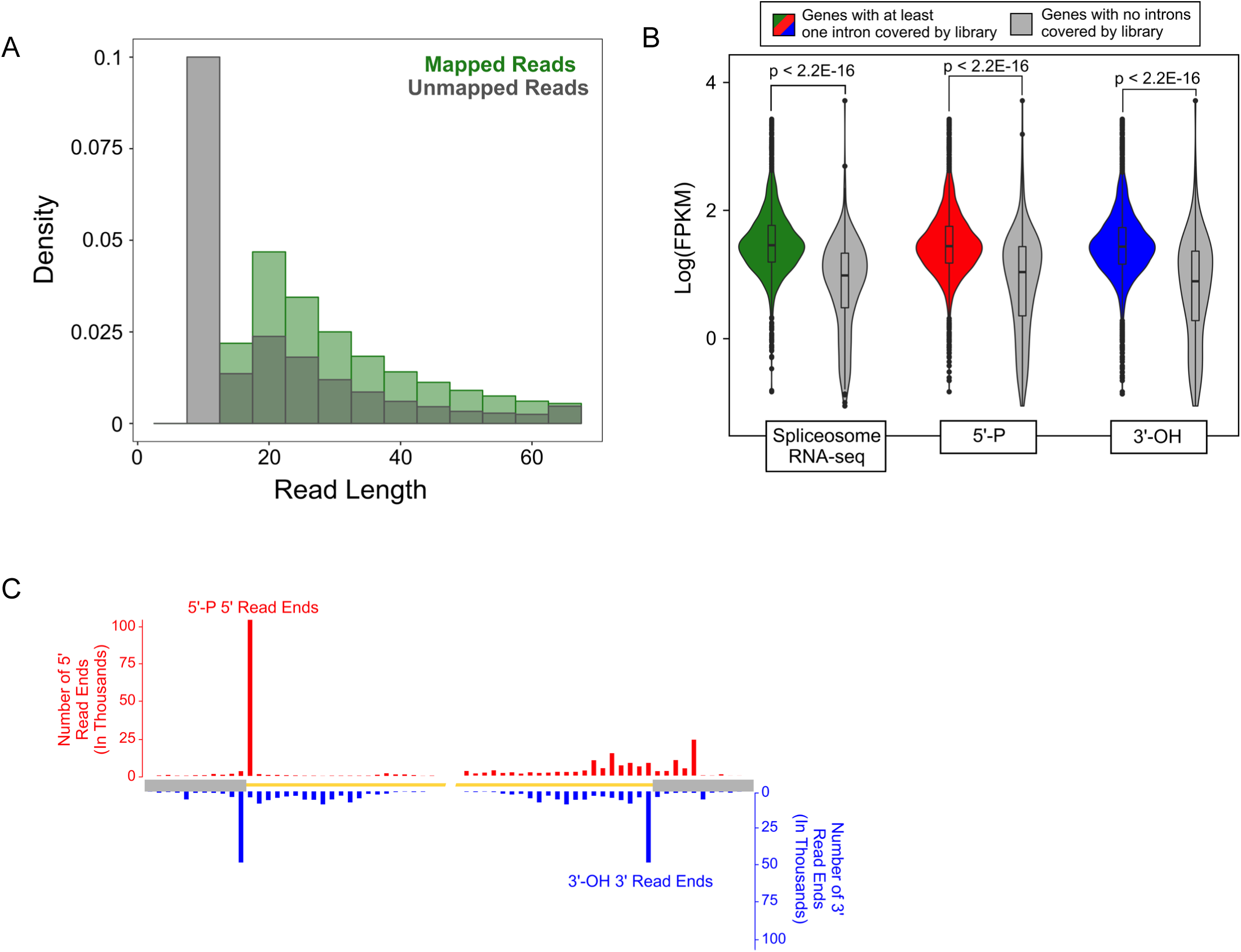
Mapped and unmapped read lengths, expression levels of intron-containing genes with or without detectable 5’ - P and 3’ - OH peaks, and 5’ - P and 3 - OH coverage on U 6 snRNA, Related to Figure 1. (A) Distribution of read lengths for mapped (green) and unmapped (gray) reads from the spliceosome RNA-Seq libraries. (B) Violin plots comparing gene expression (FPKM) of genes with introns covered by spliceosome RNA-Seq (green), 5’-P (red), or 3’-OH (blue) libraries. FPKM values were calculated from Broad polyA+ RNA-Seq libraries (ERR135906 and ERR135907; https://www.ncbi.nlm.nih.gov/pubmed/23101633) (Marguerat et al., 2012). Statistical test: Wilcoxon rank sum test. (C) Aggregation plot of all TT DBE+ 5’-P 5’ read ends (red) and 3’-OH 3’ read ends (blue) across 5’ and 3’ splice sites of the intron in U6 snRNA.

**Figure S2.**
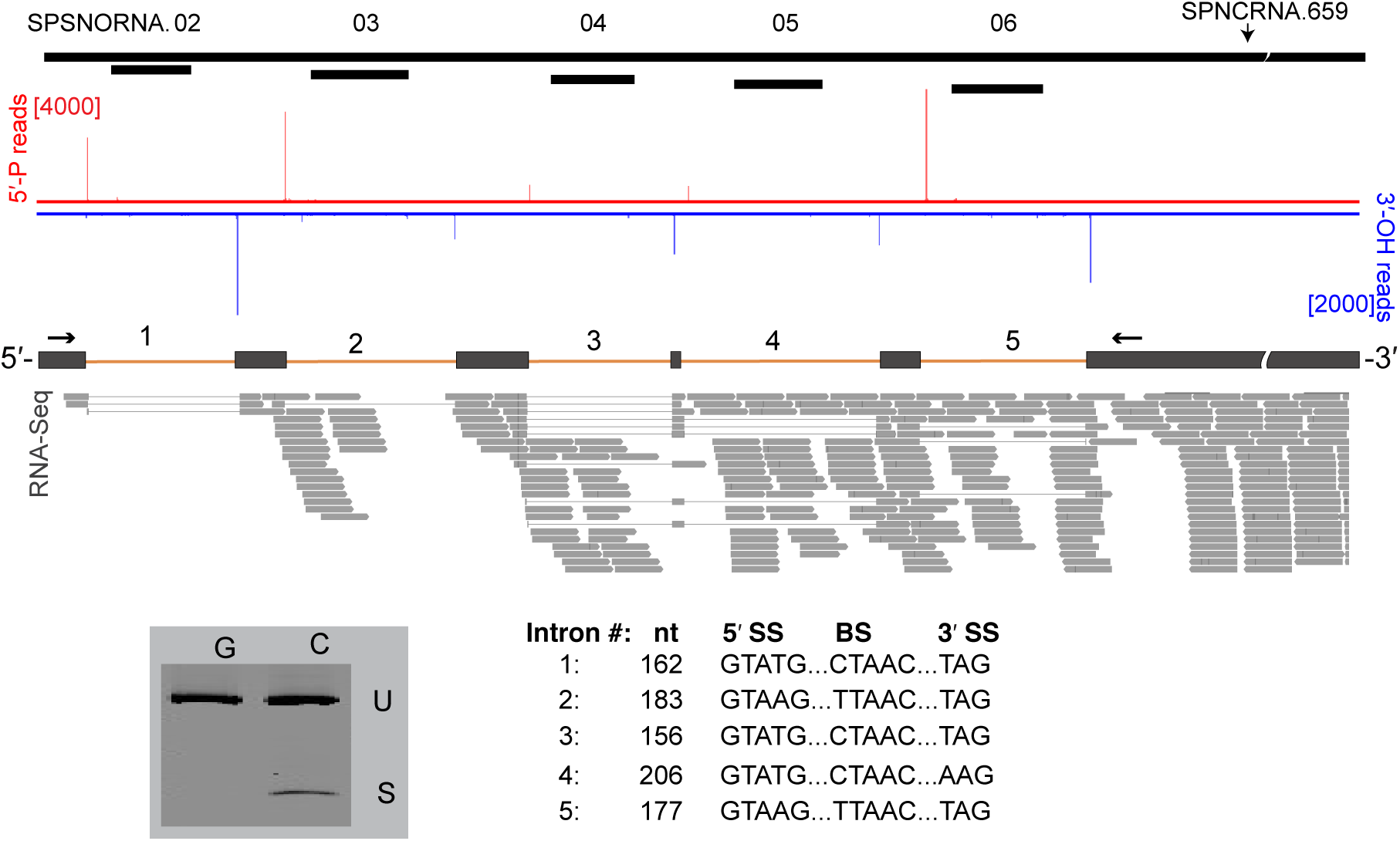
New introns in SPNCRNA. 659 revealed by 5’-P and 3’ - OH libraries, Related to Figure 2. Five new introns in snoRNA host gene SPNCRNA.659. Top track: ASM294v2.35 annotation of SPNCRNA.659 gene. Middle tracks: 5’-P (red) and 3’-OH (blue) read ends from a single biological replicate (read numbers in brackets). Bottom track: Positions of five new introns and WT polyA+ RNA-Seq reads supporting them. Bottom left: SYBR gold-stained polyacryamide gel of PCR products (U: unspliced; S: spliced) from genomic DNA (G) and cDNA (C). Bottom right: New intron lengths and their of 5’SS, BS, and 3’SS consensus sequences.

**Figure S3.**
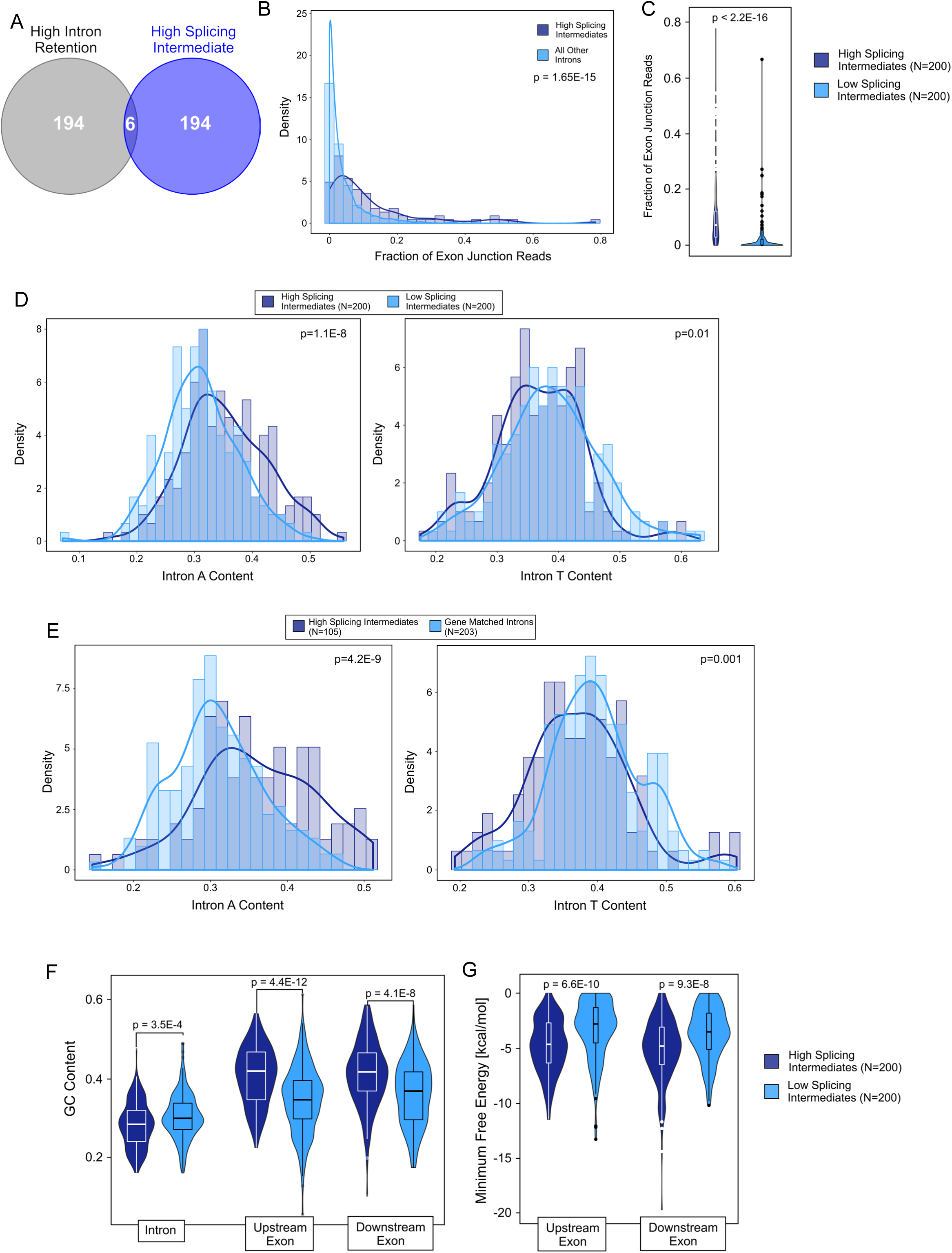
Comparison of A content, exon junction reads, GC content and minimum folding energy for high and low splicing intermediate introns, Related to Figure 3. (A) Overlap of intron sets with highest intron retention and highest splicing intermediate. (B) Histograms and density plot comparing the fraction of exon junction reads from spliceosome footprinting data between introns with high (>17%) splicing intermediate to all other introns. (P-values: Wilcoxon rank sum test). (C) Violin plots comparing of fraction of exon junction reads for the top 200 introns with highest level of splicing intermediate to the 200 introns with lowest splicing intermediate. P-values: Wilcoxon rank sum test. (D) Histograms and density plots comparing A and T content of top 200 introns with highest levels of splicing intermediate to 200 introns with lowest levels of splicing (P-values: Wilcoxon rank sum test). (E) Histograms and density plot comparing A and T content of introns with high (>17%) splicing intermediate to all other introns in the same gene (gene matched introns) (P-values: Wilcoxon rank sum test). (F) Violin plots comparing GC content of introns and exons (Upstream Exon: −1 to −40 nts from 5’SS; Downstream Exon: +1 to +40 nts from 3’SS) for sets described in B. P-values: Wilcoxon rank sum test. (G) Violin plots comparing folding free energy (RNAFold) for upstream and downstream exon sequences for sets described in B. P-values: Wilcoxon rank sum test.

**Figure S4.**
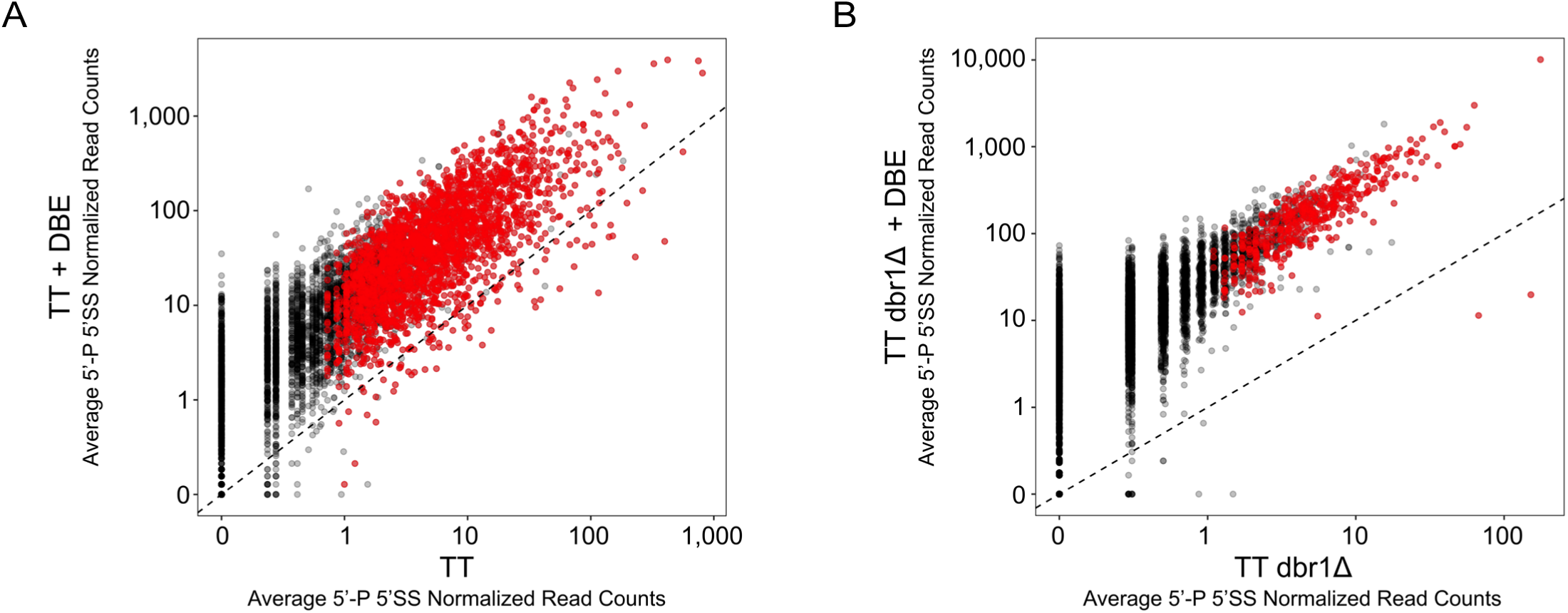
5’ - P peak correlation between DBE- and DBE+ 5’- P libraries, Related to Figure 4. (A) Scatterplot of normalized read numbers (reads per million averaged between replicates) of 5’-P reads at the 5’SS of introns between TT DBE- and DBE+ 5’-P libraries. In red (N=2,483) are introns with a significant peak in both replicates. (B) Scatterplot of normalized read numbers (reads per million averaged between replicates) of 5’-P reads at the 5’SS of introns between TT dbr1Δ DBE- and DBE+ 5’-P libraries. In red (N=472) are introns with a significant peak in both replicates.

**Figure S5.**
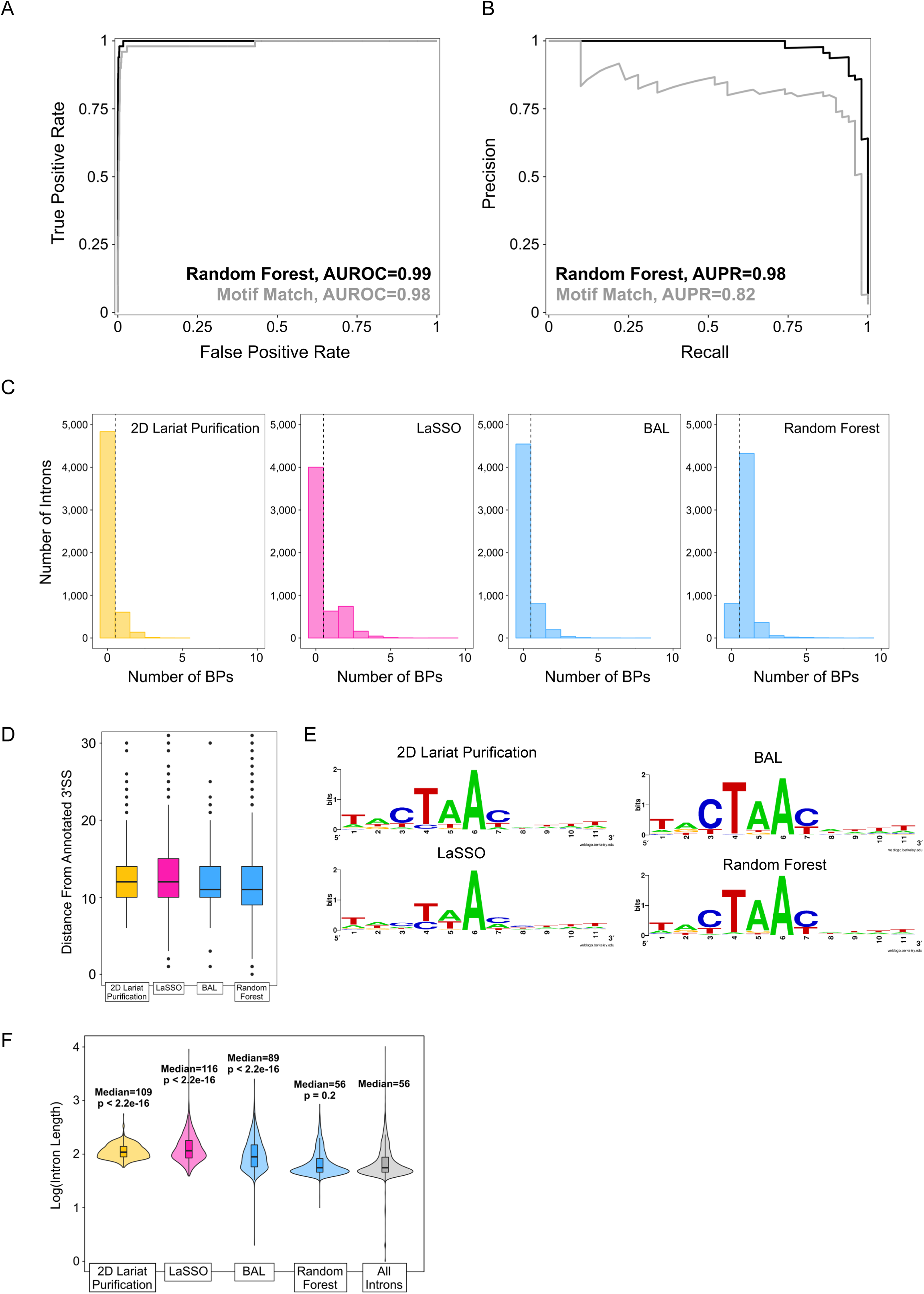
Comparison of Random Forest Method to other BP identification methods, Related to Figure 7. (A) Receiver Operator Characteristic (ROC) curve for Random Forest Model (black) and motif consensus method (gray). (B) Precision Recall (PR) curve for Random Forest Model (black) and motif consensus method (gray). (C) Histograms of the number of BPs identified per intron from 2D intron lariat purification (yellow) (Awan et al., 2013), LaSSO (pink) (Bitton et al., 2014), BAL (blue) and the Random Forest Model (blue). (D) Box plots of distance to the annotated 3’SS from BPs identified by indicated methods. (E) Sequence logos for 11 nt window centered at BPs identified by each method. (F) Violin plots of intron length for introns with BPs identified by indicated methods. All methods except for the Random Forest Model are biased toward identifying BPs in longer introns (P-values: Wilcoxon rank sum test compared to background of all introns).

## STAR ★ METHODS

### KEY RESOURCES TABLE

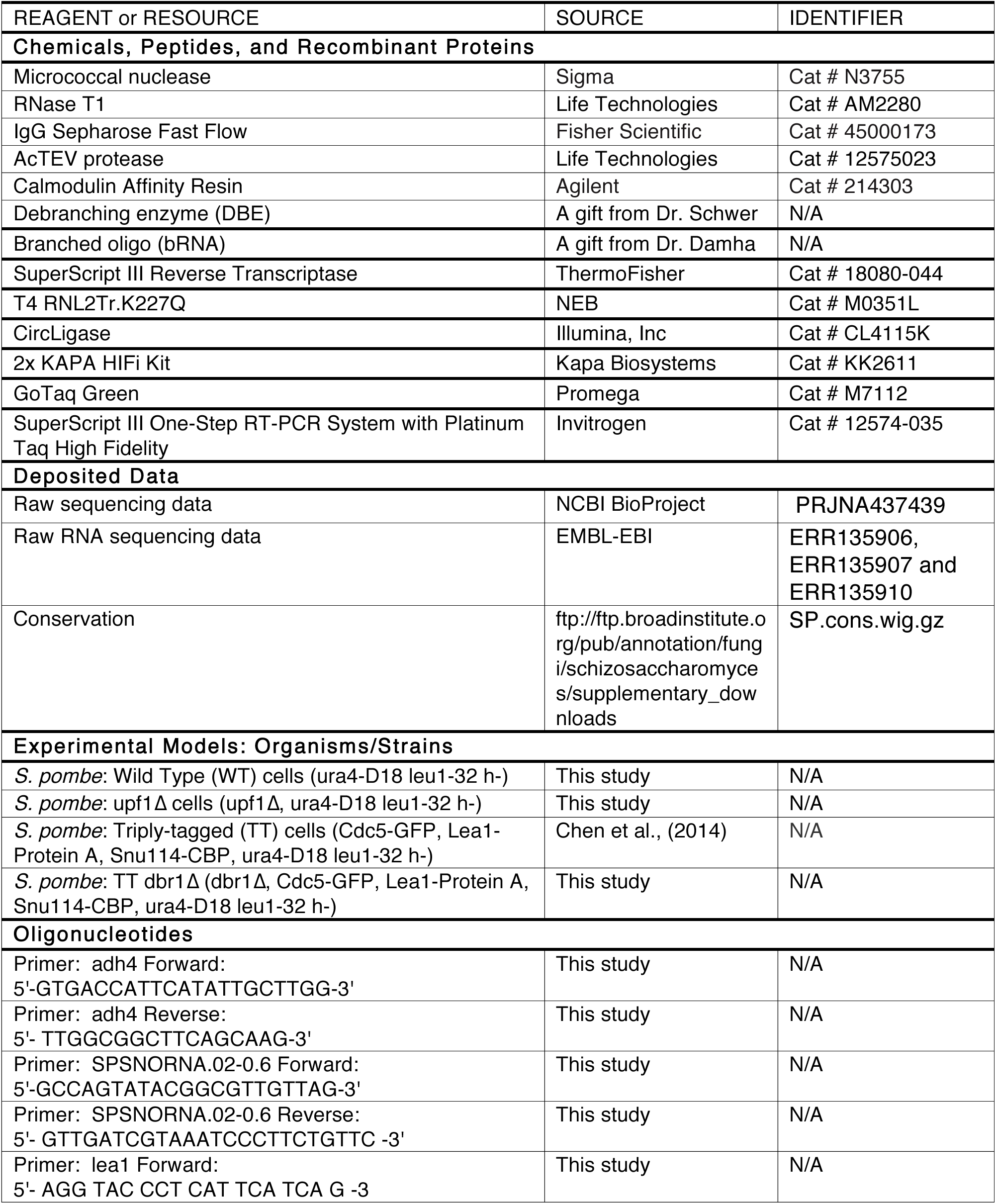
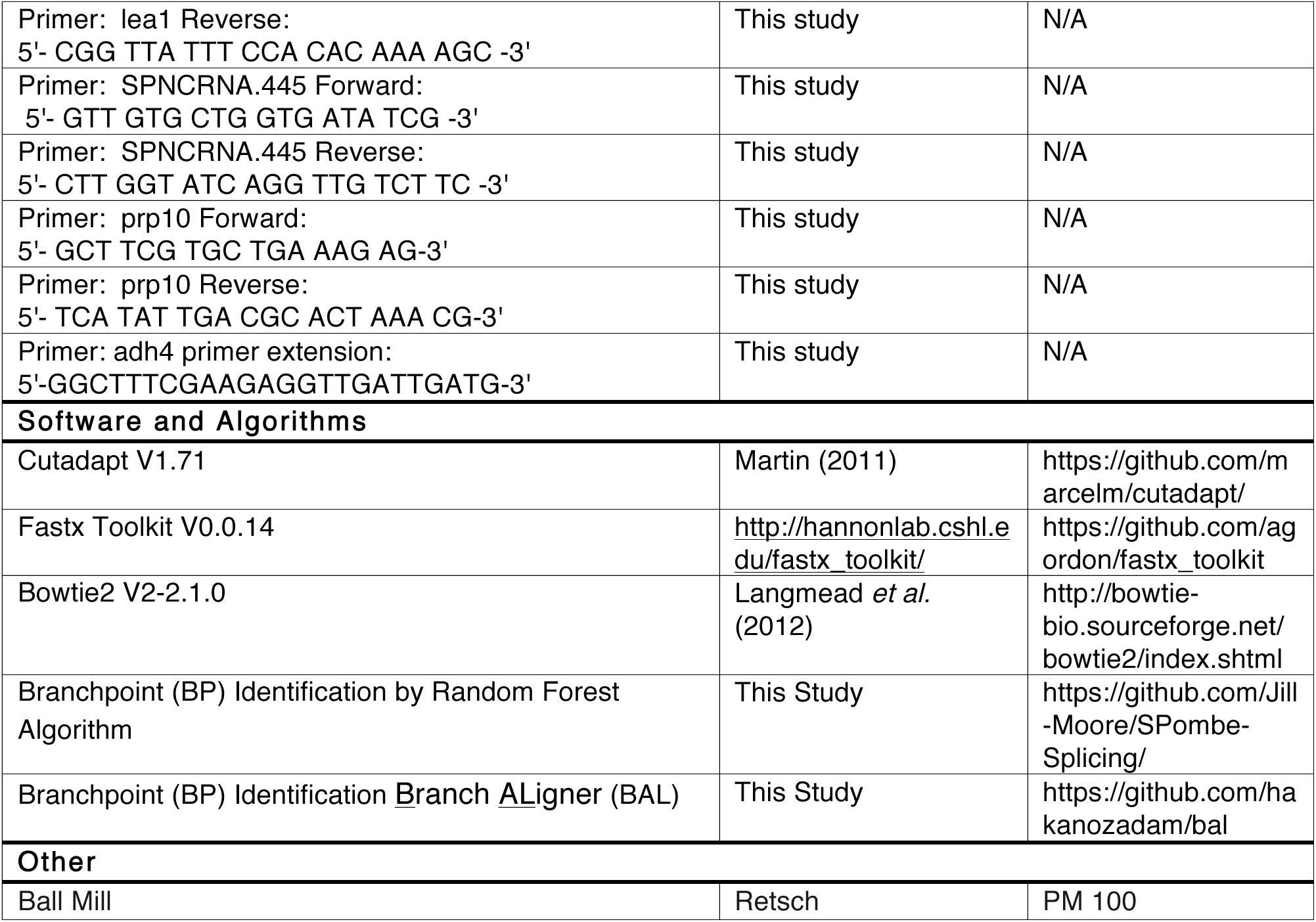

### CONTACT FOR REAGENT AND RESOURCE SHARING

Further information and requests for resources and reagents should be directed to and will be fulfilled by the Lead Contact, Melissa Moore.

### EXPERIMENTAL MODEL AND SUBJECT DETAILS

#### S. pombe strains

Strain MJM01 (herein called Triply-tagged: TT) carrying Protein A-tagged Lea1 (a U2 snRNP protein), calmodulin binding peptide (CBP)-tagged Snu114 (a U5 snRNP protein) and GFP-tagged Cdc5 (an NTC protein) were as previously described (Chen et al., 2014). Strain MJM04 (Lea1-Protein A, Snu114-CBP, Cdc5-GFP, dbr1Δ, ura4-D18leu1-32 h-) was generated by mating of MJM01 with a dbr1Δ strain from the Bioneer knockout collection (Kim et al., 2010), followed by sporulation and PCR validation of haploid derivatives (here named TT dbr1Δ).

### METHOD DETAILS

#### U2·U5·U6·NTC complex purification

Large scale (6 L initial cell culture) U2·U5·U6·NTC complex purification under High Salt (HS) conditions and MNase digestions (footprinting) were carried out essentially as previously described (Chen et al., 2014). The cell culture, cell lysis and purification details were previously published as supplementary online material to (Chen et al., 2014); they are provided again here for sake of completeness.

##### Cell Culture

Cultures (1 L) (TT cells or TT dbr1Δ cells) were grown in 5xYES in 2.8 L flasks at 30°C to OD_600_ ∼22. Cells were harvested by centrifugation at 5000 RPM 4°C for 20 minutes in 1 L centrifuge bottles. Cell pellets (∼30-35 ml from 1 L) were resuspended in an equal volume of chilled water, repelleted, resuspended in an equal volume of chilled Buffer A (10 mM K-Hepes, pH 7.9, 40 mM KCl, 1.5 mM MgCl_2_, 0.5 mM DTT, 0.5 mM PMSF, 2 mM benzamidine), and repelleted. Dry cell pellets were extruded from a 50 ml syringe directly into liquid nitrogen to form “noodles”. Frozen cell noodles were stored at −80°C until required.

##### Cell Lysis

Frozen cell noodles from 6L culture (∼200 g) were placed in a 500 mL stainless steel jar pre-chilled in liquid nitrogen. Cells were pulverized using 26-30 20 mm pre-chilled, stainless steel balls by shaking in a Planetary Ball Mill PM 100 (Retsch) for 6 × 3 min at 400 RPM. Between each cycle, jars were reimmersed in liquid nitrogen to ensure the cells remained frozen. After 6 cycles, generally 80% of cells are broken. This frozen cell powder can be stored at −80°C until required.

##### Cell Extract

All spliceosome preps described in this paper were purified under the High Salt (HS) conditions described in (Chen et al., 2014). Specifically, frozen cell powder was incubated with an equal volume of HS Buffer B (10 mM K-Hepes, pH 7.9, 300 mM NaCl, 400 mM KCl, 1.5 mM MgCl_2_, 0.5 mM DTT, 0.5 mM PMSF, 2 mM benzamidine, 10 % glycerol) with stirring at 4°C for 30 minutes to 1 hour. Extract was clarified by spinning in a preparative centrifuge at 23,000 g for 1 hour, and then the supernatant was spun in an ultracentrifuge at 100,000 g for 1 hour more. Following ultracentrifugation, three phases are generally visible: a lipid phase floating on top, a pellet of cellular debris at the bottom, and transparent a middle phase. This middle phase (S100 extract) was recovered and used for subsequent purification steps. This S100 extract was adjusted with an appropriate volume of 2 M Tris-Cl, pH 8.0, 5 M NaCl, and 10 % NP-40 to yield final concentrations equal those in to IPP 700 (10 mM Tris-Cl, pH 8.0, 700 mM NaCl, 0.1 % NP-40).

##### TAP Purification

All binding and elution steps were performed in 50 ml Econo-Columns or 0.8 × 4-cm Poly-Prep columns (Bio-Rad, Hercules, CA) as noted below. For large scale purification (∼240 ml S100 extract from 6 L initial cell culture), 20 ml of 50% slurry containing IgG Sepharose beads (Pharmacia Piscataway, NJ) is transferred to a 50 ml column and washed with 45 ml IPP 700. To promote binding of the protein A tag to the IgG beads, adjusted S100 extracts were rotated with the washed beads in sealed columns for 3 hours at 4°C. Following supernatant removal by gravity flow, beads were washed with 3 × 50 ml 2 × 50 ml IPP 700 followed by 2 × 50 ml IPP150 (10 mM Tris-Cl, pH 8.0, 150 mM NaCl, 0.1 % NP-40).

For MNase footprinting, beads were washed with 3 × 20 ml MN buffer (10 mM Tris-HCl. pH 7.4, 15 mM NaCl, 60 mM KCl, 1 mM CaCl_2_, 0.15 mM spermine and 0.5 mM spermidine). MNase digestion was carried out by adding 60U MNase (Sigma Aldrich Cat# N3755-50UN) in 10 ml MN buffer. After gentle mixing for 10 minutes at room temperature, beads were washed with TEV cleavage buffer (IPP 150 plus 0.5 mM EDTA and 1 mM DTT). For RNase T1 digests (data used in BAL section), beads were washed with 3 × 20 ml RNase T1 buffer (20 mM Tris-HCl. pH 7.6, 50 mM NaCl, 0.5 mM EDTA). RNase T1 digestion was carried out by adding 50 µL RNase T1 (Life Technologies Cat# AM2280) in 10 ml RNase T1 buffer. After gentle mixing for 15 min at room temperature, beads were washed with TEV cleavage buffer.

For TEV elution, beads were equilibrated with 2 × 50 ml TEV cleavage buffer by gravity flow. TEV cleavage was initiated by adding 8000 U AcTEV protease (Life Technologies Cat # 12575023) in 4 ml TEV cleavage buffer, followed by nutation at 16 °C for 2 hours. Eluate (∼4 ml) was recovered by gravity flow. Beads were washed with 2 ml Calmodulin Binding Buffer (CBB: 10 mM Tris-Cl, pH 8.0, 10 mM 2-mercaptoethanol, 150 mM NaCl, 1.0 mM magnesium acetate, 1 mM imidazole, 2 mM CaCl_2_, 0.1 % NP-40), which was subsequently combined with the TEV eluate.

For the calmodulin affinity step, 1.4 ml of a 50% slurry of calmodulin beads (Stratagene, La Jolla, CA) were added to a 0.8 × 4-cm Poly-Prep and washed with 10 ml CBB. The combined TEV eluate was made competent for calmodulin binding by adding 2 ml CBB and 20μl 1 M CaCl_2_ per 6 ml elutate. This solution was added to the column containing the washed calmodulin beads and rotated overnight at 4 °C. After washing by gravity flow with 30 ml CBB, bound complexes were eluted with ∼3 ml Calmodulin Elution Buffer (CEB: 10 mM Tris-Cl, pH 8.0, 10 mM 2-mercaptoethanol, 75 mM NaCl, 1 mM magnesium acetate, 1 mM imidazole, 0.01% NP-40, 6 mM EGTA). Eluate was collected in 400 μl fractions; generally ∼95% of GFP fluorescence and total protein was contained in fractions 2-4, which the peak fraction containing 1-2 mg/ml total protein.

#### RNA purification and deep sequencing library preparation

For all library preparations from purified U2·U5·U6·NTC complex, we isolated RNA from ∼40 µg complex (as measured by total protein) using hot acidic phenol, followed by chloroform extraction and ethanol precipitation. Cellular RNA was purified similarly, starting with 10 mL 1xYES cultures grown to OD_600_ = 1.5.

##### Spliceosome RNA-Seq and MNase footprinting libraries

For spliceosome RNA-Seq libraries, 250 ng total RNA from purified U2·U5·U6·NTC complex was first subjected to random hydrolysis at 94°C for 3 min in 5× first strand buffer (250 mM Tris-HCl, pH8.3, 375 mM KCl, 15 mM MgCl_2_) to generate a fragment distribution centered around 60 nts. For MNase footprinting libraries, this initial hydrolysis step was omitted because the RNA had already been fragmented will still associated with U2·U5·U6·NTC complex. Following ethanol precipitation, debranched (DBE+) samples were generated by incubating 250 ng RNA with 100 ng recombinant debranching enzyme (DBE) in 100 µL DBE buffer (50 mM Tris-HCl pH 7.0, 8 mM MnCl_2_, 2 mM MgCl_2_, 25 mM NaCl, 0.01% Triton X-100, 2.5 mM DTT, 0.15% glycerol) for 1 hour at 30°C. Reactions were stopped by adding phenol/chloroform, and the debranched RNA was ethanol precipitated. For library preparation, 250 ng of branched (DBE-) or debranched (DBE+) RNA was loaded onto a 1.5 mm thick 8% denaturing polyacrylamide gel. After excision of the region corresponding to the desired fragment length (∼10-100 nts), RNA was eluted, phenol extracted and ethanol precipitated. Recovered fragments were then treated with T4 polynucleotide kinase in the absence of ATP to remove 3’-phosphates and subjected to 3’-adaptor ligation, reverse transcription (RT) and cDNA circularization as previously described (Heyer et al., 2015). Barcodes, 3’ adaptor and RT primer sequences are shown in **Table S1**.

##### 3′-OH libraries

3’-OH library preparation was similar to spliceosome RNA-Seq library preparation except that samples were not subjected to random hydrolysis and both the size selection and 3’-dephosphorylation steps were omitted.

##### 5’-P libraries

For 5’-P library preparation, unfragmented total RNA samples from cells or purified U2·U5·U6·NTC complex were treated (DBE+) or not (DBE-) with recombinant DBE as above and then used directly for library construction employing a protocol similar to that described for making degradome libraries (Moran et al., 2014), except that 5’ adapter (aka, RNA linker) ligations were carried out for 2 hours at 25°C and ethanol precipitations were substituted for cleanup steps calling for Agencourt RNAClean XP beads. Also the RNA linker (**Table S1**) was modified to contain a region of nine random nucleotides in order to label each captured RNA fragment with a unique molecular identifier (UMI). RT and PCR primer sequences (**Table S1**) were exactly as in (Moran et al., 2014).

#### PCR confirmation of corrected, new and recursively spliced introns

Corrected introns in lea1 intron 1, prp10 intron 2, new introns in adh4, SPNCRNA.659, and recursive intron in SPNCRNA.445 gene loci were confirmed by comparing PCR products derived from genomic DNA or by reverse transcription (RT) of total cellular RNA from strains as indicated in main text. Primer sequences are in Key Resources Table. PCR from genomic DNA was performed with GoTaq Green (Promega, Cat. #M7112) following the manufacturer’s instructions. RT-PCR was performed using SuperScript III One-Step RT-PCR System with Platinum Taq High Fidelity (Invitrogen, Cat. #12574-035). All detected bands were confirmed by direct Sanger sequencing of gel-purified PCR products.

#### Primer extension assays

For primer extension analysis of adh4 mRNA isoforms, we used as input 40 µg total RNA isolated from TT cells grown at 30°C to: (1) OD_600_ = 1.5 in 1×YES medium and harvested immediately (log phase cells); (2) OD_600_ = 1.5 in 1×YES medium, then supplemented with 100 µg/mL cycloheximide and incubated for an additional 2 hours at 30°C (log phase +CHX); or (3) OD_600_ = 23 in 6×YES medium (late log glucose depletion). Samples were annealed with 1 ng 5’-^32^P labeled Primer A: 5’-GGCTTTCGAAGAGGTTGATTGATG-3’ (∼300,000 cpm) in Annealing Buffer (5 mM Tris pH 8.3, 75 mM KCl, 1 mM EDTA) by first heating for 1 minute in a boiling water bath and then incubating at 48°C for 45 minutes. Primer extension was performed with SuperScript III RT (Invitrogen) according to manufacturer’s instructions. Following ethanol precipitation, pellets were resuspended in 3.5 µL 40 µg/mL RNase A and incubated at room temp for 3 min to degrade residual RNA. Labeled extension products were then separated on an 8% denaturing polyacrylamide gel.

#### Debranching assay using labeled branched oligo

Cell extracts and debranching assays were based on previous publications (Chapman and Boeke, 1991) (Khalid, 2005). Briefly, 50 ml of TT or TT dbr1Δ cells grown in 1xYES to mid-log (OD at 600 nm ∼1.0) were harvested by centrifugation, washed once with cold water and resuspended in 400 ul cold buffer A (1 M sorbitol, 50 mM Tris pH 8.0, 10 mM MgCl_2_, 3 mM DTT) plus 300 ul glass beads. Cells were broken by vigorous shaking on a small ball mill for 4 min at 4°C. After addition of 20 ul Buffer B (300 mM HEPES pH 7.8, 1.4 M KCl, 30 mM MgCl2), extracts were incubated on ice for 20 min and then cell debris removed by centrifugation (13,000 rpm for 5 min at 4°C in a bench top centrifuge). After transferring the supernatant to a 3 ml tube (Beckman Cat. # 362305), a second high speed spin (58,000 rpm for 60 min at 4°C in a Beckman TLA 110 rotor) yielded clarified extract, which was divided into 100 ul aliquots, quick frozen in liquid N_2_ and stored at −80°C.

The branched oligo (25 pmol bRNA-AK86 ((Clark et al., 2016); sequence shown in **Figure 4C**) was 3’-end-labeled with T4 RNA ligase (NEB Cat. # M0204S) and 20 µL ^32^P-pCp (Perkin Elmer # NEG019A250UC) as previously described (Nilsen, 2014). Preliminary labeling experiments revealed that the ^32^P-pCp as supplied by the manufacturer contained residual γ-^32^P-ATP and T4 polynucleotide kinase activity, which led to inadvertent 5’-end labeling of bRNA-AK86 and circle formation during the overnight T4 RNA ligase incubation. To prevent these undesired side reactions, residual T4 polynucleotide kinase was inactivated by incubating ^32^P-pCp at 65°C for 20 min prior to addition to the T4 RNA ligase reaction. Ligation reactions were carried out overnight at 4°C and then unincorporated ^32^P-pCp removed by spinning the reaction through an AM10070 NucAway Spin Column.

For detection of debranching activity, control reactions (20 µl) contained Buffer A (50 mM Tris–HCl pH 7.0, 8 mM MnCl_2_, 2.5 mM DTT, 25 mM NaCl, 2mM MgCl_2_, 0.01% Triton X-100, 0.15% glycerol) alone (**Figure 4C**, lane 1) or plus 100 ng purified *S. cerevisae* DBE (Khalid et al., 2005) (lane 2). Cell extract reactions (20 µl) contained Buffer B (50 mM Tris–HCl pH 7.0, 8 mM MnCl_2_, 2.5 mM DTT, 0.01% Triton X-100, 0.15% glycerol), 700 ng purified total *S. pombe* cellular RNA, 5-7 µl TT (lane 3) or TT dbr1Δ (lane 4) cell extract (300 µg total protein), and 3 fmol 3’-^32^P-pCp-labeled bRNA-AK86. All reactions were incubated at 30°C for 1 hour and then an aliquot resolved by 20% denaturing PAGE.

#### Deep sequencing and read mapping

##### Spliceosome-seq Libraries

Sequencing specifics (e.g., sequencing platform used, bar codes, read lengths, etc.) and read count statistics are given in **Table S1**. All libraries will be deposited in the NCBI short read archive upon manuscript acceptance. Scripts for all bioinformatics analyses available at: https://github.com/weng-lab/Jill_Moore/tree/master/Moore/Scripts

Using Cutadapt (V1.7.1) (Martin, 2011), we trimmed 5’ barcode and 3’ adaptor sequences from read pairs. Then using custom python scripts (available on GitHub) or fastx collapse (fastx toolkit V0.0.14), we collapsed reads into non-redundant species and removed the randommers from each species. We then aligned reads to the S. pombe genome using Bowtie2 (V2-2.1.0) and default parameters. We selected all uniquely mapping, concordant read pairs for additional analysis. We then mapped the remaining reads to exon-exon junction sequences, selecting those that uniquely map and combining them with the uniquely mapping genomic reads.

##### WT RNA-Seq Libraries

ERR135906, ERR135907 and ERR135910 RNA-Seq datasets were downloaded from the SRA database and were mapped to the *S. pombe* genome using HISAT2 (V2-2.0.0-beta) with default parameters.

### QUANTIFICATION AND STATISTICAL ANALYSIS

#### Binomial test for significant 5’-P and 3’ - OH peaks

##### Annotated Introns

We tested for significant 5’-P and 3’-OH read-end peaks at annotated intron ends using the binomial test. At annotated 5’SSs, we counted the number of 5’-P reads that started exactly at the 5’SS compared to those starting at any position in a 25 nt downstream window. At annotated 3’SSs, we counted the number of 3’-OH reads that ended exactly at the 3’SS compared to those ending in a 25 nt upstream window. To quantify the amount of splicing intermediate, we also analyzed the number of 3’-OH reads ending one nt upstream of the 5’SS and compared it to the number of reads ending in a 25 nt window centered at the 5’SS. In each case we tested for significance using a binomial distribution:

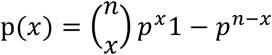

where n is the number of reads in the window, p is 1/length of window, and x is the number of reads at the splice site or −1 position. We applied an FDR correction and selected all peaks with FDR < 0.01.

##### Genome Wide

To identify significant peaks 5’-P and 3’-OH elsewhere in the genome we counted the number of 5’-P reads starting at and 3’-OH reads ending at every genomic location not previously identified as a 5’SS or 3’SS and compared this number to the number of reads ending in a 25 nt window centered around that position. We calculated the significance of each peak using the aforementioned binomial test, applied an FDR correction, and reported all peaks with FDR < 0.01.

#### Calculating Intron Retention

For each intron, we calculated the percent of intron retention using RNA-Seq data as follows:

A = Number of exon junction reads spanning the intron

B = Number of reads that cross the 5’SS. Reads must extend at least 5 nts on either side of the splice site.

C = Number of reads that cross the 3’SS. Reads must extend at least 5 nts on either side of the splice site.

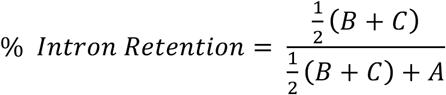

#### BAL Algorithm

We first mapped libraries against the *S. pombe* genome using the HiSat exon junction aligner to identify genome- and exon junction-mapping reads (Kim et al., 2015). Unmapped reads were then locally aligned using Bowtie2 (Langmead and Salzberg, 2012) against the first 20 nucleotides downstream of every 5’SS. Mapping to a particular 5’SS was confirmed if the 3’ remainder (dark green region in **Figure 6B**) mapped to the intron sequence immediately downstream. The 5’ remainder (purple region in **Figure 6B**) was then mapped to the downstream intron to identify the BP. To increase depth by allowing for inclusion of shorter reads, we then generated a BP sequence reference list by concatenating the first 100 nts of each intron with the 100 nts upstream of each identified BP. All non-mapped reads were then mapped to this reference using Bowtie2 in the end-to-end alignment mode. B reads were defined as those with read length >22 and mapping quality >5; BP confirmation required a >70% mismatch rate at the BP among all reads mapping to that reference sequence.

#### Random Forest

The Random Forrest Method relies on reads terminating at or crossing BPs in 3’-OH DBE-libraries. For some introns, however, we did not initially observe such reads from the initial genome-mapping pipeline because the BP was so close to the 3’SS (< 15 nts) that these short reads did not map to the genome. To capture these additional short reads that cannot map uniquely when compared to the entire genome, we remapped previously unmapped reads using only introns as our reference. This yielded an additional 1.3 million (increase of ∼10%) mapped reads for the 3’ OH DBE-libraries.

To create our training, validation, and testing sets, we used 300 BPs from 291 introns that were identified by both BAL and 2D lariat purification (Awan et al., 2013) (**Table S4B**). The identified BPs were our positives and all other “A” nts within these 291 introns were our negatives. We included 192, 49, and 50 introns in the training, validation, and testing sets, respectively (which represented 200, 50 and 51 BPs, respectively).

For training, we included eight features in our Random Forest model:

1. Distance from BP to 3’SS
2. % of total intron reads starting at 3’SS and stopping at BP
3. % of total intron reads starting at 3’SS and stopping one nucleotide downstream of BP
4. % of total intron reads starting one nucleotide 5’ to BP
5. Mismatch % (mutation and/or indel) at BP for reads starting at 3’SS (i.e., from lariat product)
6. Mismatch % (mutation and/or indel) at BP for reads starting in following exon (i.e. from lariat intermediate)
7. Nucleotide immediately 5’ to BP (A, C, G or T)
8. Nucleotide immediately 3’ to BP (A, C, G or T)

We ran each model ten times, generating a Random Forest with 100 trees each time. For prediction, we averaged the score across all ten iterations. We then applied the model genome-wide and reported all predictions with probability > 0.5.

To assess the validity of our Random Forest Model, we compared its performance to an algorithm that relied solely on the BP consensus motif. For each BP we calculated its log-odds score match to the consensus motif using FIMO and ranked predictions from high to low score. To evaluate each method, we calculated the area under the receiver operator characteristic (ROC) and precision recall (PR) curves using the ROCR and PROCR R packages. We also calculated precision and recall using FIMO with Q-value cutoffs of 0.01 and 0.05.

### DATA AND SOFTWARE AVAILABILITY

The bioproject accession number for the raw data files for the RNA sequencing analysis reported in the paper is PRJNA437439.

Cutadapt V1.71 (Martin, 2011) is available at GitHub (https://github.com/marcelm/cutadapt/), Fastx Toolkit V0.0.14 (http://hannonlab.cshl.edu/fastx_toolkit/) is available at GitHub (https://github.com/agordon/fastx_toolkit), Bowtie2 V2-2.1.0 (Langmead *et al*. (2012) is available at Bowtie (http://bowtie-bio.sourceforge.net/bowtie2/index.shtml), BP Identification by Random Forest Algorithm is available at GitHub (https://github.com/Jill-Moore/SPombe-Splicing/), BAL is availbe at GitHub (https://github.com/hakanozadam/bal)

## Supplemental table legends

**Table S1: Libraries used in this study, Related to All Figures**

A. Library specifics and raw read numbers

B. Transcript class read mapping statistics for spliceosome RNA-Seq and footprinting libraries (Round I).

C. Transcript class read mapping statistics for spliceosome RNA-Seq and footprinting libraries (Round II).

D. Transcript class read mapping statistics for 5’-P libraries.

E. Transcript class read mapping statistics for 3’-OH libraries.

**Table S2: New and previously misannotated introns, Related to Figure 2**

A. Previously misannotated introns in CDS genes (16)

B. New intron-containing transcripts (18)

C. New introns in noncoding RNA genes (119)

D. New introns in CDS genes (80)

**Table S3: High splicing intermediate introns, Related to Figure 3**

A. Introns with high splicing intermediate

B. Gene ontology (Top 100)

C. Gene ontology (Top 150)

D. Gene ontology (Top 200)

**Table S4: Identified branch points, Related to Figure 6 and 7**

A. BPs identified by BAL

B. BPs included in Training, Validation, and Test Sets for Random Forest Model

C. BPs identified by Random Forest Model

